# Integrating Phenomic Selection Using Single-Kernel Near-Infrared Spectroscopy and Genomic Selection for Corn Breeding Improvement

**DOI:** 10.1101/2024.09.07.611678

**Authors:** Rafaela P. Graciano, Marco Antônio Peixoto, Kristen A. Leach, Noriko Suzuki, Jeff Gustin, A. Mark Settles, Paul R. Armstrong, Márcio F. R. Resende

## Abstract

Phenomic Selection (PS) is a cost-effective method proposed for predicting complex traits and enhancing genetic gain in breeding programs. The statistical procedures are similar to those utilized in genomic selection (GS) models, but molecular markers data are replaced with phenomic data, such as near-infrared spectroscopy (NIRS). However, the use of NIRS applied to PS typically utilized destructive sampling or collected data after the establishment of selection experiments in the field. Here, we explored the application of PS using non-destructive, single-kernel NIRS in a sweet corn breeding program, focusing on predicting future, unobserved field-based traits of economic importance, including ear and vegetative traits. Three models were employed on a diversity panel: G-BLUP and P-BLUP models, which used relationship matrices based on SNP and NIRS data, and a combined model. The genomic relationship matrices were evaluated with varying numbers of SNPs. Additionally, the P-BLUP model trained on the diversity panel was used to select doubled haploid (DH) lines for germination before planting, with predictions validated using observed data. The findings indicate that PS generated good predictive ability (e.g., 0.46 for plant height) and effectively distinguished between high and low germination rates in untested DH lines. Although GS generally outperformed PS, the model combining both information yielded the highest predictive ability, with considerably higher accuracies than GS when low marker densities were used. This study highlights the potential of NIRS both to achieve genetic gain where GS may not be feasible and to maintain/improve accuracy with SNP-based information while reducing genotyping costs.

**Key message:** Phenomic selection using whole seeds is a promising alternative to improve genetic gain and complement genomic selection in corn breeding. Models that combine genomic and phenomic data maximize the predictive ability.

## 1 Introduction

Genetic progress in breeding has traditionally relied on phenotypic evaluations conducted through field trials. Over recent decades, molecular markers have shown promise in assisting the selection process. The use of markers together with phenotypic data from historical individuals is used to build a predictive model. This model is then applied to predict the phenotype of a new set of genotyped individuals in what is called genomic selection (GS). This technique has gained popularity due to its potential to increase genetic gain (Meuwissen et al. 2001; Bernardo and Yu 2007; Heslot et al. 2015; Xu et al. 2020). Although there have been advancements in genotyping techniques, the cost of genotyping individuals can still limit its application, especially in small breeding programs.

Recently, phenomic selection (PS) has been proposed as an affordable and effective tool for predicting complex traits (Rincent et al. 2018). This approach uses statistical models similar to GS, but instead of using molecular marker data, such as SNPs, it uses multivariate phenotyping methods, like near-infrared (NIR) spectroscopy data. NIRS measures the emission or reflection of light from a sample across a specific wavelength range. Since chemical bonds absorb light at different wavelengths, the NIRS data is related to the biochemical properties of a sample (Robert et al. 2022b). In the traditional use of NIRS, a model is trained to predict a trait (e.g. starch content) for the same sample where the NIRS data is collected. Partial least squares regression is the method most utilized to properly weight the reflectance or absorbance wavelengths that better explain the content and composition of the trait being predicted (Beć et al. 2021). This approach is routinely used in many breeding programs to, for example, phenotype kernel composition in grains, like oil, starch, and protein content (Jiang et al. 2007; Sundaram et al. 2009; Chadalavada et al. 2022), lignin content in wood (Yeh et al. 2004), and feed quality in forages (Decruyenaere et al. 2009; Samadi et al. 2020).

Phenomic selection differs from the classical use of NIRS because it aims to predict traits relevant to the breeding program but not necessarily related to the biochemical properties of the sample scanned by the NIRS. These may involve predictions of different environments or quantitative traits for which the genes affecting the trait are expressed in different tissues from the ones where the NIR spectra are collected. It is hypothesized that the predictive power of such an approach can come from i) the ability of a multivariate phenotype to estimate genetic relationships, which can, in turn, be used to predict other phenotypes of interest, or ii) the ability to exploit the endophenotypes, which represents intermediate molecular layers between the genome and the phenotype, by associating them with the target trait (Posada et al. 2009; Rincent et al. 2018; Zhu et al. 2021, 2022; Robert et al. 2022a). Therefore, NIRS could predict traits unrelated to the analyzed sample (Robert et al., 2022b). Studies have shown the potential of PS in different scenarios, with some obtaining prediction ability comparable to or higher than GS (Rincent et al. 2018; Krause et al. 2019; Cuevas et al. 2019; Lane et al. 2020; Zhu et al. 2021, 2022; Robert et al. 2022a; Weiß et al. 2022; Brault et al. 2022; Adunola et al. 2024).

Some of these studies explore PS with hyperspectral imaging, allowing for a nondestructive analysis. However, hyperspectral imaging often requires collecting data after the selection experiments have already been established in the field and/or at a time point that is close to when the target trait is expressed (Aguate et al. 2017; Krause et al. 2019; Parmley et al. 2019; Galán et al. 2020). This allows for selection before harvest while avoiding expensive phenotyping, but it still requires waiting periods comparable to those of phenotypic selection. It is essential to assess the efficiency of PS with data collected from young tissue. It is possible to achieve this when using NIRS, especially with seed data. Some literature examples involve grinding the seeds (Rincent et al. 2018; Weiß et al. 2022; Adunola et al. 2024), making the process destructive and less efficient, and limiting its application in cases where the number of seeds is limited in the early selection stages.

Single-kernel NIR (skNIR) spectroscopy, which collects data on individual kernels, offers advantages like low cost, high speed, and non-destructiveness, although sensitivity can be a limitation (Hacisalihoglu and Armstrong 2023). It has proven useful for assessing grain sample variability, sorting specific traits, and detecting rare attributes. For example, in corn, skNIR has been shown to sort kernel composition mutants (Spielbauer et al. 2009), predict kernel composition differences in transgenics (Lappe et al. 2018), and differentiate haploid from diploid seeds (Gustin et al. 2020). Additionally, skNIR is effective in phenotyping kernel composition before planting, thereby enhancing trait selection (P. R. Armstrong 2006; Gustin et al. 2013; Hacisalihoglu et al. 2020, 2022; Fan et al. 2022). Using whole kernels to predict complex traits before planting could improve throughput and reduce cycle time, achieving similar reductions as GS but at a lower cost. Additionally, the use of skNIR spectroscopy for PS before planting has not been explored and needs further investigation to determine feasibility.

In this study, we explored PS in sweet corn (*Zea mays L*.) using skNIR. Sweet corn is a popular vegetable in the United States and Canada, and breeding programs focus on hybrid development to address producer and consumer needs (Lertrat and Pulam 2007; Hu et al. 2021; Peixoto et al. 2024a). Our aim was to investigate the use of this technology for predicting economically important sweet corn traits, such as ear and vegetative traits. We assessed the effectiveness of PS using a diverse panel of inbred lines and validated the process using doubled haploid (DH) elite lines. Our objectives were to: (1) investigate the effect of the source of NIRS data in phenomic prediction, (2) compare the predictive ability of PS, GS, and a multi-matrix model using NIRS and molecular markers, and (3) assess the feasibility of scanning and performing prediction before planting.

## 2 Materials and methods

### 2.1 Experimental design and phenotypic data

Model calibration and evaluation were performed using a diversity panel of 693 sweet corn inbred lines from the Sweet Corn Coordinated Agricultural Project (SweetCAP). This panel consists of tropical and temperate adapted materials, such as open-pollinated landraces, historically important inbred lines, and modern elite inbred cultivars, as well as elite lines from the University of Wisconsin-Madison Sweet Corn Breeding Program and the University of Florida Sweet Corn Breeding Program. Both mutations used in sweet corn production, *sugary1* (*su1*) and *shrunken2* (*sh2*) genes, are represented in the diversity panel. These genes impact endosperm starch biosynthesis and are responsible for increased sugar content in sweet corn relative to field corn (Hu et al. 2021). The lines were grown at the University of Florida Plant Science Research and Education Unit in Citra, Florida, USA, over three years (2019, 2021, and 2022). A randomized resolvable incomplete block design was used with six blocks and two replicates in 2019, and 30 blocks and three replicates were used in subsequent years. Checks were used in all years with at least one check per block.

A total of 24 traits from two different categories were measured. Vegetative-related traits included GER: germination; DTP: days to pollination; DS: days to silking; PH: plant height; EH: ear height, FLH: flag leaf height; TE: tassel extension TN: tiller number; LA: leaf angle; TBN: tassel branch number; AC: anther color; SC: silk color; NES: number of ear shoots. Ear-related traits included: EL: ear length. EW: ear width; TPF: tip fill; HAP: husk appearance; KRN: kernel row number; SL: shank length; SOL: solidity; TP: taper; CUR: curvature; COV: convexity; and KRF: kernel row fill. Table S1 provides a complete list of all measured traits. For 2021 and 2022 seasons, some ear-related traits (EL, EW, COV, CUR, KRF, SOL, SL, TP, and TPF) were measured with EarCV, a computer vision tool that allows measuring quality traits relevant to sweet corn from photographs (Gonzalez et al. 2022).

For model validation, we evaluated doubled haploids (DHs) lines produced from 15 F_1_ families. Each F1 used to create these doubled haploid families was generated by crossing 27 elite sweet corn inbreeds from the sweet corn breeding programs at the University of Florida (24 inbreds) and the University of Wisconsin (3 inbreds). Seeds from 522 DHs were produced in Chile and scanned using a skNIR machine. The model calibrated with the diversity panel set was used to predict germination rates. Based on model predictions, nine genotypes with high predicted germination rate (above 70%) and nine genotypes with low predicted germination rate (below 55%) were selected (Table S2 shows the predicted values for the selected individuals and their rankings among all 522 individuals). These seeds were planted in Citra, Florida, in 2024 in a completely randomized design with three repetitions, considering the two groups (high and low germination). As an additional validation, four DHs out of the 18 predicted high and low germinating lines described above (2 with low and 2 with high germination) were used to evaluate the effect of seed source on the model predictions. New seeds for these genotypes were produced in Idaho in 2023. The additional seed source of the four genotypes was planted in Citra in the same experiment described above.

### 2.2 Phenotypic analysis

In the diversity panel, we implemented individual analysis for each trait and in each environment to compute the best linear unbiased estimation (BLUE), which was later used for the second stage of the analysis. The statistical mixed model used was as follows:

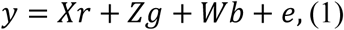

where y is the vector of phenotypic values; *r* is the vector of fixed replicate and checks inside blocks effects, added to the overall mean; *g* is the vector of fixed genotypic effects; *b* is the vector of random block effects (considered random), where 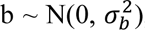, being 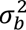 the block variance; and *e* is the vector of residuals, 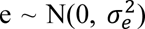, where 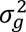 is the residual variance. X, Z, and W represent the incidence matrices for *r*, *g*, and *b*, respectively.

The estimation of variance components was based on the model from equation 1, but considering the genotypic term as random, such that 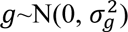, where 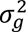 is the genotypic variance. Broad-sense heritability was derived from the variance components estimates as the ratio between genotypic and phenotypic variance (Piepho and Möhring 2007), as follows:

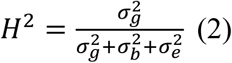

The significance of the genotypic effect was tested using the likelihood ratio test (LRT), with a chi-square statistic with 1 degree of freedom and a 5% probability of Type I error (Wilks 1938).

### 2.3 NIRS data

Two custom-built grain analyzers were used to collect skNIR data in the Sweet Corn Breeding and Genomics laboratory at the University of Florida (Armstrong, 2006). Both instruments use a model CDI 256L-1.7T1 with a 256-element InGaAs photodiode array that spans the spectral range from 906 to 1679 nm with a step size of 1 nm (Control Development Inc., South Bend, Ind.). General operation involves dropping kernels into an illuminated glass tube (light tube), while automatically triggering the spectrometer to collect the spectrum as the kernel travels down the tube. A bifurcated optical fiber looking down both ends of the tube is used to collect light. A slightly angled pathway allows a clear view of the kernel by the fiber within the light tube. One version requires manual feeding of individual kernels. The second version, hereafter referred to as the skNIR-sorter, uses a commercial vacuum corn planter singulator to feed kernels to the light tube and a solenoid sorting system to sort kernels based on a measured constituent level. Both instruments are controlled by custom electronic hardware and PC-based software to provide a broad range of user control. This includes applying user developed calibration models, selection of sorting thresholds, storing individual spectra, real-time statistics of measured values, and detecting poor quality spectra for exclusion. The skNIR-sorter instrument measures and sorts kernels at a little more than two kernels per second.

The skNIR data was collected in three datasets. Two datasets were obtained from the diversity panel with kernels harvested in Florida, one in 2019 and the other in 2020. The 2019 dataset was composed of kernels produced in the Everglades Research and Education Unit at Belle Glade, FL (14% of genotypes) and kernels produced at the Plant Science Research and Education Unit in Citra, FL (86% of genotypes). In 2020, the entire dataset was produced at the Plant Science Research and Education Unit in Citra, FL. Up to four ears were harvested at the mature stage for each genotype. After drying, a subset of 25 single intact kernels per genotype was run through the skNIR instrument. Spectral data was collected from both skNIR instruments for a subset of 256 lines from the 2020 seed production year, and the performance of the two instruments was similar based on predictability from model calibrations for each instrument using the same kernels (Fig. S1). PS models were validated with skNIR-sorter data from the set of 522 doubled-haploid lines. In this dataset, up to 25 kernels per genotype were scanned, each being scanned three times.

Any spectra with values that deviated by more than three standard deviations from the mean of the individual wavelength per genotype were considered outliers and removed. The remaining spectra, which had already been automatically normalized by the skNIR software to a mean center of 1, were then averaged per genotype. It is known that external factors independent of sample composition, such as temperature, can influence the measurement (Robert et al. 2022a). We tested additional preprocessing methods in the diversity panel set to address these external effects that can bias spectra comparison. The following methods were evaluated: Standard Normal Variate (SNV), Multiplicative Scatter Correction (MSC), Detrend (DT) and first derivative. The SNV, MSC, and DT pre-processing were applied to the averaged mean-centered spectra. SNV centers and scales the spectra, while MSC normalizes by aligning each spectrum with an ideal reference one, free of scattering effects (in our case, the ideal reference was the average of all spectra retained after outlier filtering). Both SNV and MSC are normalization methods that can correct light scatter effects. The DT applies an SNV transformation followed by a second-degree polynomial regression (Barnes et al. 1989). Additionally, the Savitzky-Golay filter was used to compute the first derivative from normalized spectra produced from SNV (SNV-1der) and MSC (MSC-1der) (Savitzky and Golay 1964). The DT and the Savitzky-Golay filter reduce noise and correct baseline shifts. For the diversity panel set, pre-processing was applied to spectra separately for each kernel source year (2019 and 2020), as well as an average across the two years.

The pretreated skNIR data (outcome from each one of the five preprocessing analyses) were used to obtain the phenomic relationship matrix (P) between pairs of individuals, following:

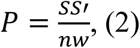

where S is the matrix of pretreated skNIR spectra, centered and scaled for each wavelength with dimensions *ng* x *nw*(where *ng* is the number of individuals and *nw* is the number of wavelengths). Two additional analyses were performed on the skNIR spectra from the diversity set: a principal components analysis (PCA) and an estimation of variance components. The PCA was conducted using both skNIR spectroscopy data and marker data (described below). In estimating variance components, each wavelength from the transformed skNIR data was treated as a separate response variable, incorporating a fixed effect of year in which kernels were harvested and a random effect for genotype by year interaction to estimate the variance components for each wavelength.

### 2.4 Genotypic data

The genotypic data of the diversity set were obtained via whole genome resequencing. Whole genome sequences were aligned to that Ia453-*sh2* genome (Hu et al., 2021) using BWA-MEM. From the total of variants called (Colantonio, 2023) an initial subset of 200k markers was randomly sampled using vcftools (Danecek et al. 2011). SNPs with a minor allele frequency of less than 1% and missing data greater than 30% were removed from the data set to ensure data quality. After filtering, a final set of 128,202 SNPs was used to estimate the additive genomic relationship matrix (G) between lines, following the VanRaden formula (VanRaden 2008).

To evaluate cases with lower marker coverage, we randomly selected 500 markers from the initial pool of 200k. Markers were added to the subset size, ensuring each subsequent subset contained all markers from the previous level. This process resulted in 6 new subsets with 500, 1500, 5000, 10,000, 50,000, and 100,000 markers. The procedure was repeated five times, with each subset using the same filtering for minor allele frequency and missing data mentioned before. The G matrix was calculated for each subset and repetition leading to subsets with averages of 321, 957, 3,200, 6,382, 32,040, and 64,112 SNPs, respectively.

### 2.5 Prediction models

To predict the breeding values in the second stage of the analyses, we used a linear mixed model, utilizing both single-trait (ST) and multi-trait (MT) approaches for the diversity panel set. The genomic best linear unbiased prediction (GBLUP) (VanRaden 2008) follows:

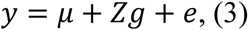

Where *y* is the vector of BLUEs values for all traits predicted from the first stage of the analyses, *μ* is the overall mean, *g* is the additive genetic effects for all traits, assumed to follow a normal distribution, 𝑔 ∼𝑀𝑉𝑁(0, 𝛴_𝑔_⊗𝐺), where *G* represents the genomic relationship matrix and 𝛴_𝑔_ the genetic variance-covariance matrix across traits. Z is the incidence matrix for g, and e is the vector of residuals 𝑒∼𝑀𝑉𝑁(0, 𝛴_𝑒_⊗𝐼), where 𝛴_𝑒_ residual variance-covariance matrices across traits, and I is an identity matrix. We employed an unstructured form for the genetic 𝛴_𝑔_ and residual 𝛴_𝑒_ variance-covariance matrices, allowing for heterogeneity of variances and the consideration of specific genetic correlations among traits.

The same model described for the genomic prediction was used for phenomic prediction. Therefore, the genomic relationship matrix (*G*) was replaced by the phenomic relationship matrix (*P*) in a P-BLUP approach. Additionally, we fitted a model incorporating both genomic and phenomic information as independent random effects.

SkNIR data was collected from kernels harvested in 2019 and 2020 in the diversity panel set. These kernels and additional ones from other sources were then planted to collect phenotypic traits across three years (2019, 2021, and 2022) (Figure 1). This allowed us to investigate the effect of kernel source on the ability of skNIR to predict the traits from 2019, 2021, and 2022. However, there was some overlap in the seed sources used for phenotypic evaluation and skNIR measurement. To ensure a fair comparison between the phenomic and genomic prediction models, we restricted the analysis to include only genotypes present in the genomic and skNIR data for each year (Table S3 provides information on the number of individuals for each analysis).

**Fig. 1.**
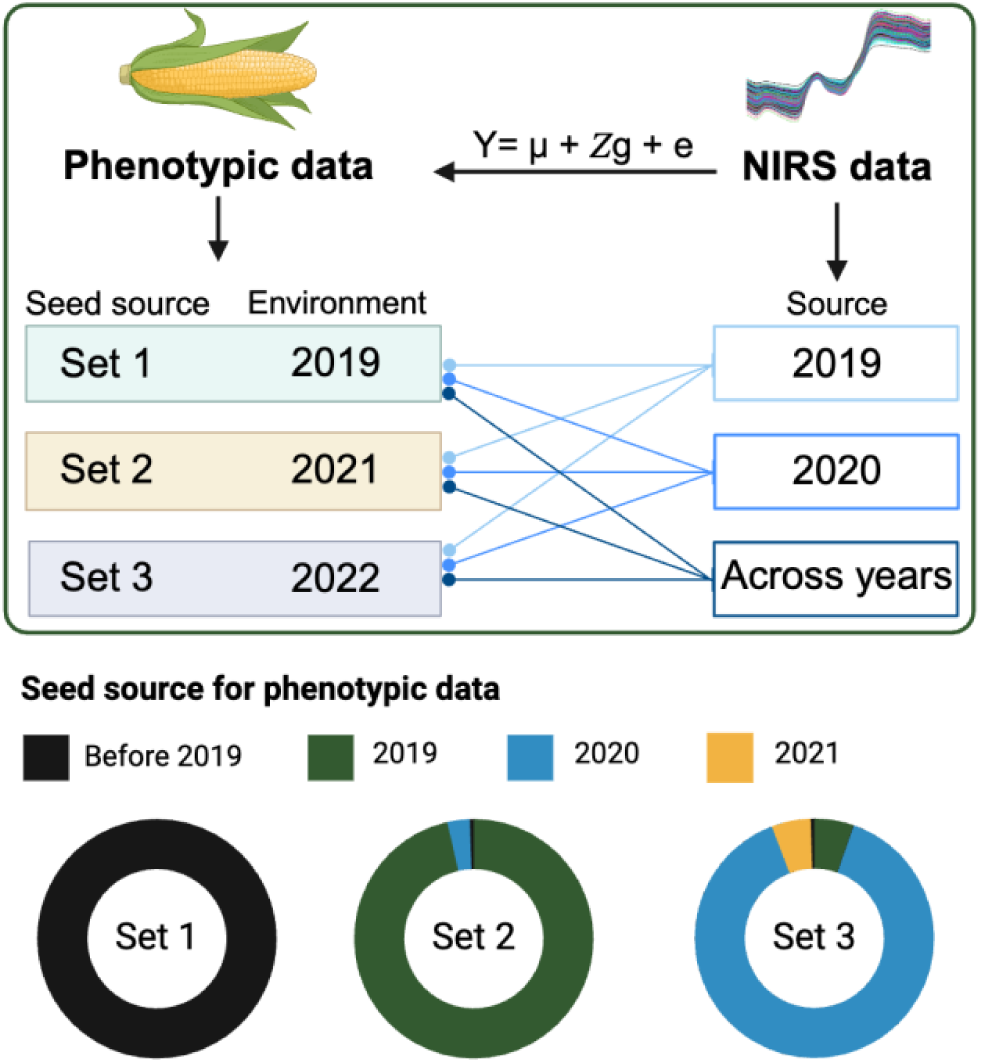
Phenomic prediction scenarios. In the diversity panel set, NIRS data was collected from kernels harvested in 2019 and 2020, while phenotypic traits for the same population were measured in 2019, 2021, and 2022. The genotypes planted for phenotypic evaluation used seeds produced in different years, with seed source sets indicating the proportion of seeds used for each year. The “before 2019” seeds came from the effort to select the diversity panel from sweet corn germplasm, with diverse sources dating from 1988 to 2018. The lines originating from the NIRS data and terminating with a circle on the phenotypic data indicate the NIRS data source used to predict the phenotypic traits of that specific year. Created with BioRender.com

Additionally, partial least squares (PLS) regression was implemented in the pls package (Mevik and Wehrens 2007). The number of components used in each PLS model was selected through 10-fold cross-validation based on the root mean squared error prediction (RMSEP) curve using the one-sigma heuristic approach. In cases where the one-sigma approach resulted in 0 components, we used 1.

The phenomic selection model used for model validation using the DHs population was calibrated using the germination rates from 2022 and skNIR data (average across data collected from kernels sourced from 2019 and 2020) from the diversity panel set. This calibrated model was then used to predict the germination rates of untested DHs using the skNIR-sorter dataset. The predictions were evaluated as described above (see *Experimental design and phenotypic data*).

### 2.6 Model evaluation

To evaluate model performance for the diversity panel set in the second stage, we performed 20 repetitions of a fivefold cross-validation. In each repetition, the individuals were randomly assigned to 5 subsets, whereas a training population (4 parts) was used to predict the testing population (1 part). The process was repeated five times to ensure every subset was used once as the testing set. Then, a Pearson correlation was calculated between the BLUE values and the estimated breeding values. We also measured the mean squared error (MSE), given by the averaged squared deviation from estimated breeding values and BLUEs. The Pearson correlation and the MSE were averaged over the 20 repetitions, providing the final predictive ability and error measures.

### 2.7 Computational implementation

The phenotypic analyses (first stage) were carried out in the R package ASREML 4.0 (Gilmour et al. 2015), and the following prediction models (second stage) were implemented in the BGLR package (Pérez and de los Campos 2014; Pérez-Rodríguez and de los Campos 2022). The Bayesian models used 10000 iterations, a burn-in of 1000, and a sampling interval (thin) of 10, totaling 900 iterations. To obtain the additive relationship matrix (*G*), we used the R package AGHmatrix (Amadeu et al. 2016, 2023), and for all the pretreatments on the spectra, we used the R package prospectr (Stevens and Ramirez-Lopez 2014). All the codes and the data used in the analyses are available at https://github.com/Resende-Lab/Graciano_skNIR_Phenomic_Seleciton.

## 3 Results

This study evaluated a diversity panel of 693 sweet corn inbred lines over three years (2019, 2021, and 2022). Analyzing the phenotypic data (Table S1 contains a summary of trait information), the LRT indicates that the genotypic effect was significant for all traits. The heritability exhibited variations, ranging from 0.12 (CUR in 2022) to 0.85 (SC in 2021). The traits evaluated in this study presented a range of pairwise phenotypic correlations ranging from −0.95 to 0.95 (Fig. S2).

We observed that different traits had different correlation patterns with skNIR spectra, ranging from −0.55 to 0.55 (Fig. 2A and Figs. S4 to S7). Vegetative traits tended to have higher correlations than ear traits, and these patterns were generally consistent across the years. Although the magnitude of the correlation varied by year, the wavelengths showing correlation were consistent between years. In 2021 and 2022, GER showed the highest correlation with specific wavelengths throughout the spectrum (with a magnitude of approximately 0.5), followed by PH (magnitude of around 0.35) (Fig. S4). However, in 2019, the correlation with germination was weaker, reaching a maximum magnitude of 0.25, while plant height maintained similar values to the other years, having, on average, the highest correlation in 2019. We also observed variation in the spectral profile for the genotypes in the panel (Fig. S3A and S3B). Upon partitioning the variance components along the SNV-1der processed spectrum, genotypic variance always accounted for more than 20%, some around 80% (Fig. S3C). On average, the proportion of genetic by environment variance was low.

**Fig. 2.**
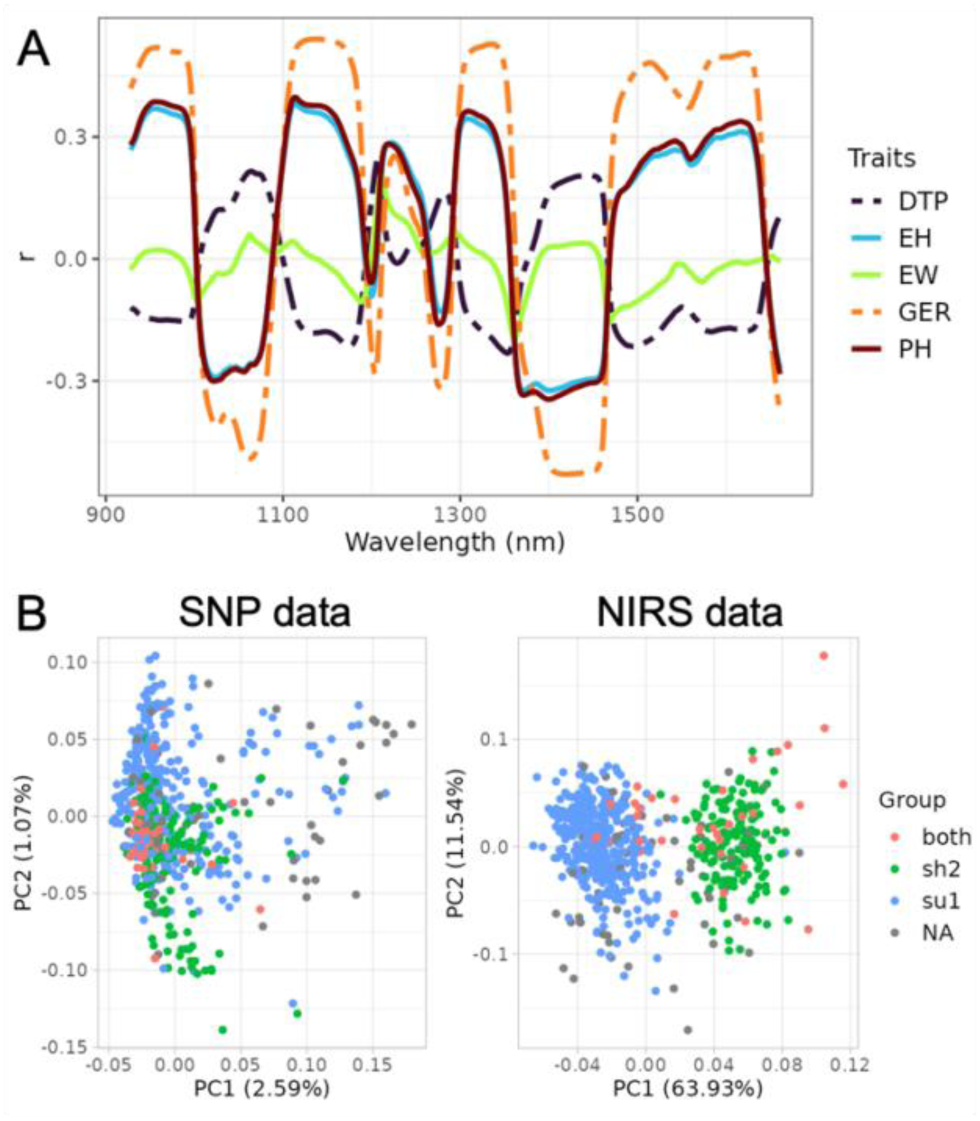
NIRS data analysis for the diversity panel. (A) shows the correlation between each wavelength and the traits from 2021 for days to pollination (DTP), ear height (EH), ear width (EW), germination (GER), plant height (PH). (B) Principal component analysis (PCA) shows the amount of variance explained by the first two components, PC1 and PC2, based on marker data (left) and NIRS data averaged from 2019 and 2020. The group indicates if the genotypes have the *sugary1 (su1)* and/or *shrunken2 (sh2)* mutation genes

PCA using the molecular markers did not delineate distinct clusters, but PC1 of the SNV-1der data skNIR analysis separated two clusters of accessions based on *su1* and *sh2* kernel phenotypes (Fig. 2B). When comparing the relationship matrices, the relationship estimated from SNV-1der skNIR data from 2019 and 2020 showed a high correlation of 0.87. In contrast, the correlations between relationship matrices derived from markers and those derived from skNIR spectra were low, around 0.13.

### 3.1 Phenomic selection demonstrated potential for accurate prediction

Using the P-BLUP approach for phenomic selection in the diversity panel, we tested five preprocessing methods for skNIR data collected from kernel produced in different years, i.e., 2019, 2020, and an average of the two years. Based on the prediction ability and MSE, it was observed that these factors affected the results, and there was some variation depending on the trait (Tables S4 and S5). The overall best method for preprocessing the spectra was the SNV-1der, which was selected for all downstream analyses.

The prediction ability of phenomic selection alone varied across different traits and years tested, with values ranging from −0.10 (TN in 2021) to 0.57 (GER in 2021) (Fig. 3 and Table S4). When looking at the average predictive ability for each of the 24 traits across all years and skNIR datasets, only 11 traits had a predictive ability lower than 0.20. In comparison, six traits had a predictive ability higher than 0.35. Among the phenotypic years assessed, GER demonstrated the highest prediction ability in 2021 and 2022 (0.56 and 0.55, respectively), but it was lower in 2019, around 0.23. The PH trait also showed good predictive ability for all years (around 0.43). The traits AC and SC color also had relatively high prediction accuracies, ranging from 0.38 to 0.44. Ear-related traits had low predictive ability, around 0.12, except EW and KRN in 2019 (0.32 and 0.31, respectively).

**Fig. 3.**
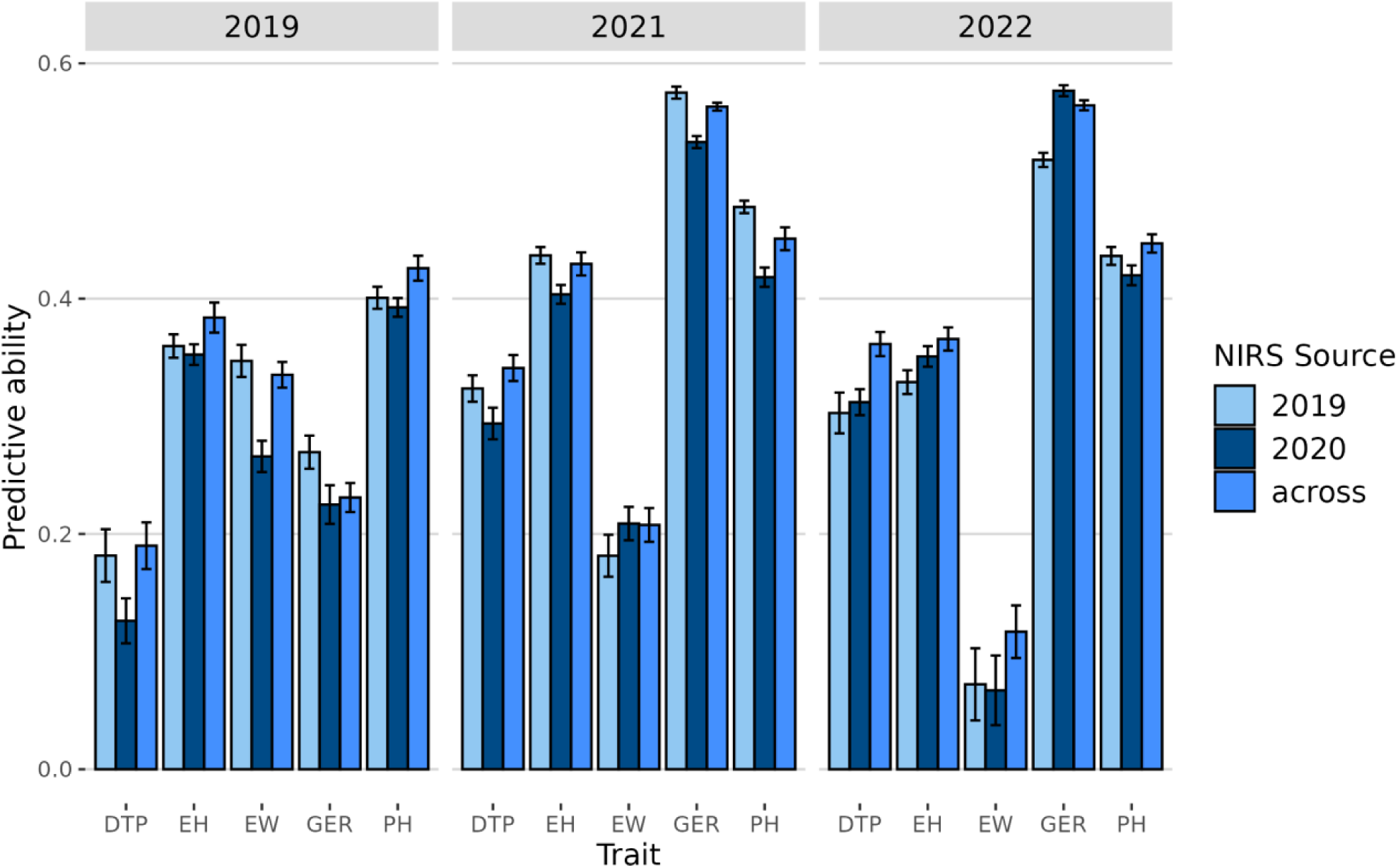
Predictive ability of phenomic selection (PS) using NIRS data collected in different years, 2019, 2020, and an average across the two years (across). The traits evaluated over three years (2019, 2021, and 2022) were days to pollination (DTP), ear height (EH), ear width (EW), germination (GER), and plant height (PH). Error bars are standard errors of means from 20 repetitions

We also used PLS regression for phenomic prediction in addition to the P-BLUP approach. Both methods generated predictions on an entry basis with skNIR spectra averaged within the genotype. The results showed that, generally, P-BLUP produces comparable or higher prediction ability than using the PLS methods (Fig. 4). Overall, BLUP was around 31% better than PLS. The change in prediction ability when moving from PLS to BLUP ranged from –0.003 in the case of BN to +0.16 in the case of HUK (Table S4).

**Fig. 4.**
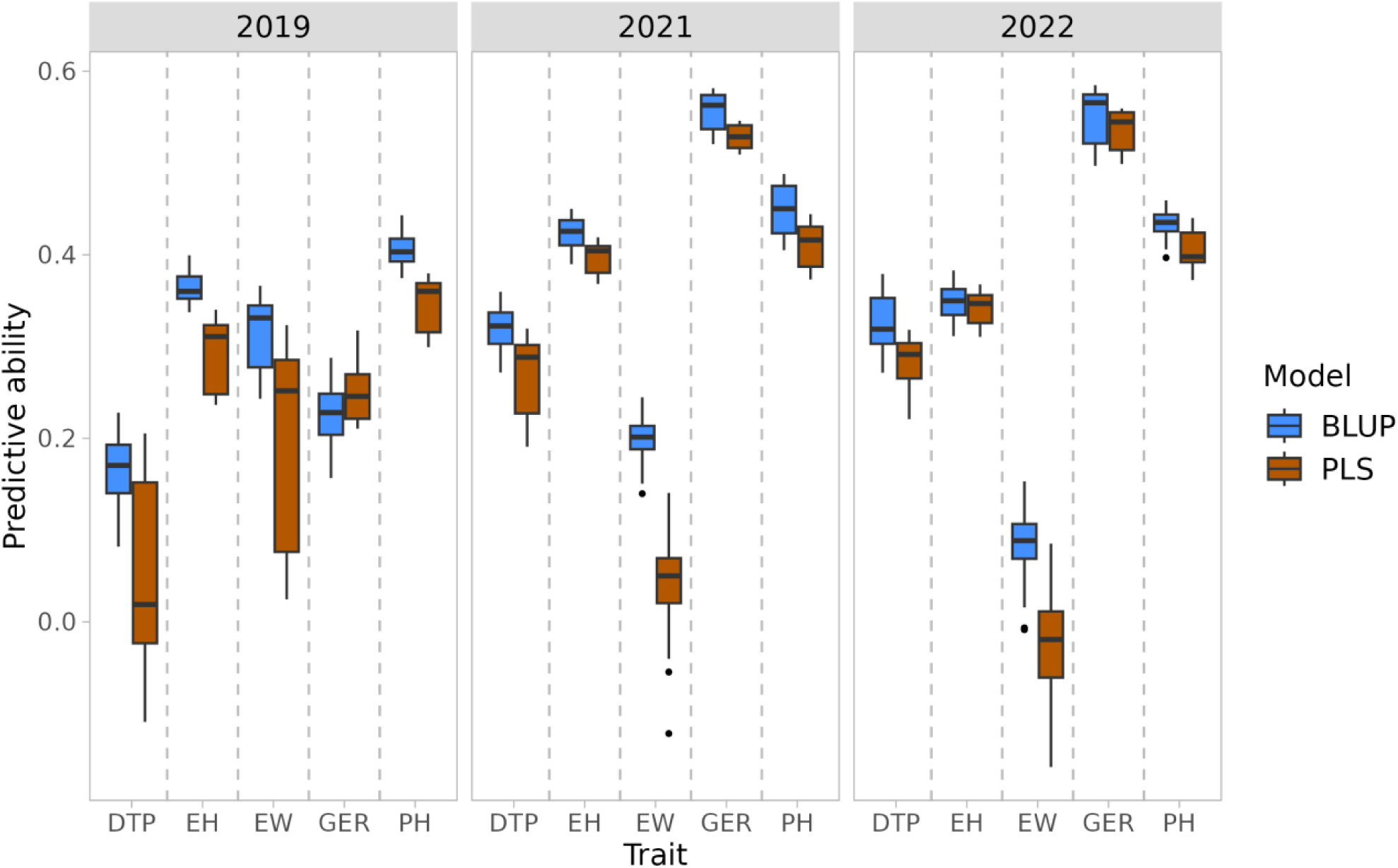
Predictive ability in the diversity panel set using NIRS data to fit two different models: PLS and NIRS-BLUP (as a phenomic selection (PS) approach). The traits evaluated over the three years (2019, 2021, and 2022) were days to pollination (DTP), ear height (EH), ear width (EW), germination (GER), and plant height (PH). Phenomic data comprised NIRS spectra processed using the Standard Normal Variate first derivative preprocessing method. The results include predictions using skNIR data collected from kernels produced in different years (2019, 2020, and the average across years)

### 3.2 NIRS data and seed source had a small effect on the prediction ability

When analyzing the results of the P-BLUP approach for skNIR data in different years, the best results were often achieved using the skNIR data averaged from two kernel sources (Fig. 3 and Table S4). However, this varied depending on the trait or the predicted environment (*i.e*., the year the traits were assessed). For example, in 2022, the best results for GER were obtained using skNIR data from the 2020 seed source, while the best results for DTP were achieved using the average across years. When predicting GER for 2021, the best result was obtained using skNIR data from the 2019 kernel source, whereas for 2022, it was based on skNIR data from the 2020 kernel source (Fig. 3). Despite these variations, the predictive values were generally similar among modeling approaches for a specific trait, with the variation in predictive ability within the phenotypic years usually being less than 0.1.

We also observe that differences in seed source used for phenotypic evaluation may affect the prediction ability of certain traits. For example, germination showed higher predictive ability when the seeds planted for phenotypic evaluation and scanned by the skNIR came from the same year (Fig. 3). The prediction ability for GER evaluated in 2021 was 0.04 higher when the skNIR spectra were collected from the 2019 seed source than the 2020, and most seeds used for planting came from the 2019 seed source. Similarly, for 2022, where most seeds used were from the 2020 seed source, skNIR spectra from the 2020 seed source was 0.06 higher than skNIR data from 2019. However, not all traits follow this pattern. For instance, using skNIR data from the 2019 seed source consistently resulted in the best values for plant height, outperforming skNIR data from the 2020 seed source by 0.06 in 2021 and by 0.02 in 2022.

Additionally, we conducted a supplemental analysis to account for the source seed effect, ensuring that the seeds scanned for skNIR data came from a different year than those planted for phenotypic evaluation. This revealed that the impact on value varied based on the trait (Fig. S8). For instance, for GER in 2022, there was a small reduction in prediction ability when there was no overlap in seed source (around 0.02), but not in 2021. Overall, the values were similar.

### 3.3 Combining phenomic and genomic data can improve prediction ability

In addition to the P-BLUP model, we evaluated two other BLUP models using marker data: genomic selection (G-BLUP) and a model that combined both phenomic and genomic information (G+P). The prediction ability values varied from 0.12 (G-BLUP for SOL in 2022) to 0.65 (G+P for DTP in 2021) (Fig. 5 and table S4). Overall, the G-BLUP model showed higher prediction ability than P-BLUP by 0.19 in 2019 and 0.16 in 2021 and 2022. However, the G+P model achieved the highest prediction ability, resulting in a 0.02 increase across all years and all traits compared to the G-BLUP model. For the G-BLUP model, the traits with the highest prediction ability differed from P-BLUP, with the DTP trait in 2021 (0.63) and the EH trait in 2019 and 2021 (0.63 and 0.62, respectively). The pattern observed in prediction ability was similarly reflected in the MSE results (Table S5).

**Fig. 5.**
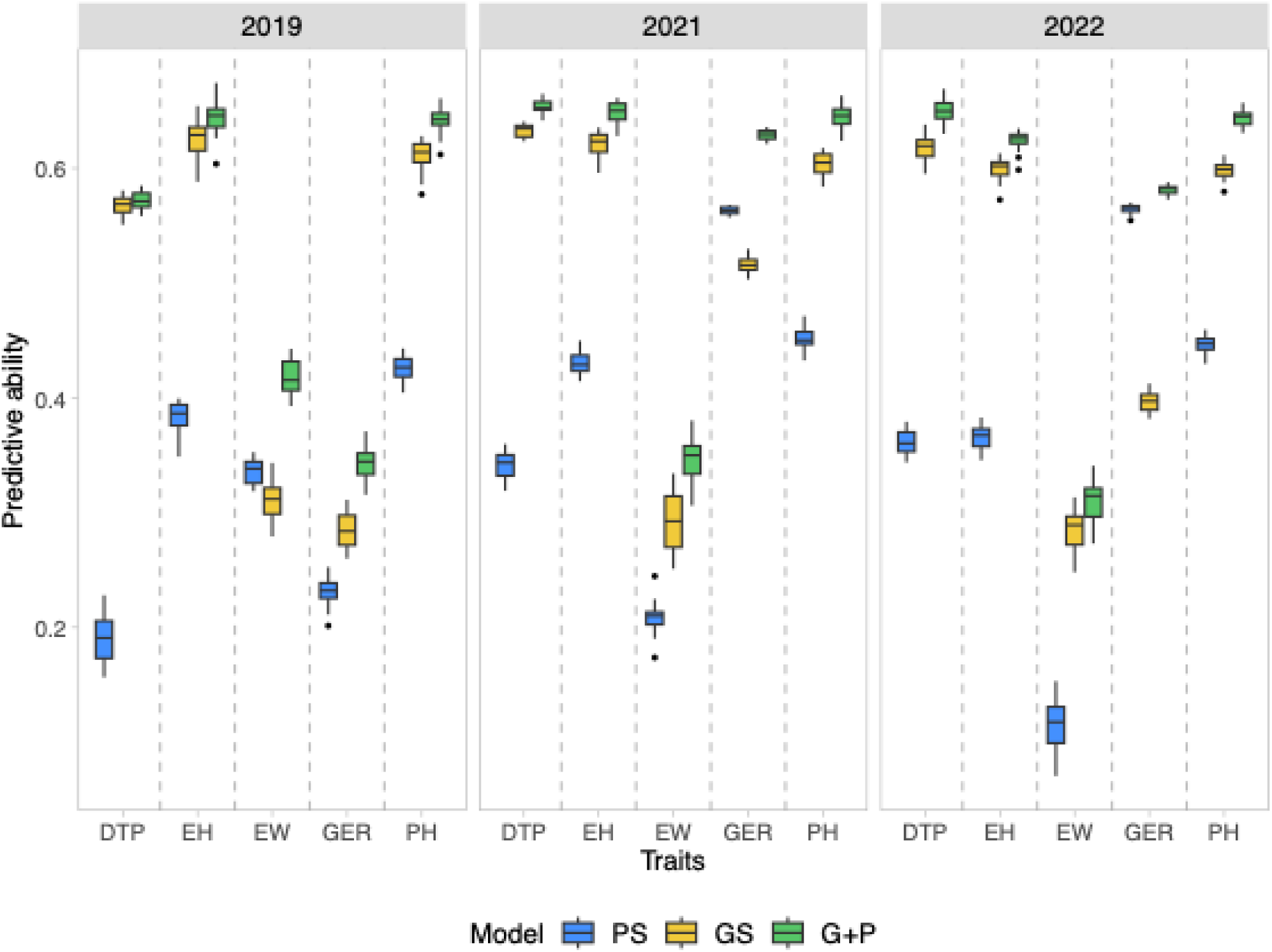
Predictive ability of single-trait approaches across genomic selection (GS), phenomic selection (PS), and a combined approach using both genomic and phenomic data (G+P) for the diversity panel set. The phenomic model used NIRS spectra averages from data collected in 2019 and 2020, processed using the Standard Normal Variate first derivative preprocessing method. Days to pollination (DTP), ear height (EH), ear width (EW), germination (GER), and plant height (PH)

The multi-trait approach did not lead to overall improvements in results. The mean predictive ability was generally comparable to or lower than that in the single-trait model (Table S4). Interestingly, there are some exceptions to the multi-trait approach. For example, the P-BLUP model performed better than the G-BLUP model for PH in 2022, a deviation from the single-trait analysis.

Additionally, we explored the G-BLUP prediction ability with different numbers of markers. Fig. 6 shows the results for seven different marker sets (averaging 321, 957, 3,200, 6,382, 32,040, and 64,112 markers) when predicting traits in 2021. The relative performance of P-BLUP to the series of G-BLUP models with a range of markers varied depending on the trait being analyzed. For instance, in the case of GER, the P-BLUP model prediction is higher even when all the markers are used in the G-BLUP model. In contrast, for DTP and PH, the G-BLUP model shows better or similar prediction ability even with only 321 markers (G-BLUP: 0.49 for DTP and 0.44 for PH, compared to P-BLUP: 0.34 for DTP and 0.45 for PH). For EW, the G-BLUP model using all the markers outperforms the P-BLUP model (0.3 for G-BLUP and 0.21 for P-BLUP). However, with fewer than 3,200 markers, the P-BLUP model performs better (e.g., 0.15 for G-BLUP with 957 markers). Prediction ability generally improves with the G+P model, which indicates that combining NIRS with genomic data increases prediction ability compared to using G-BLUP or P-BLUP models alone. This leads to accuracies higher than 0.5 for GER, PH, and DTP traits using only 321 markers. Results were similar for other years (Fig. S9).

**Fig. 6.**
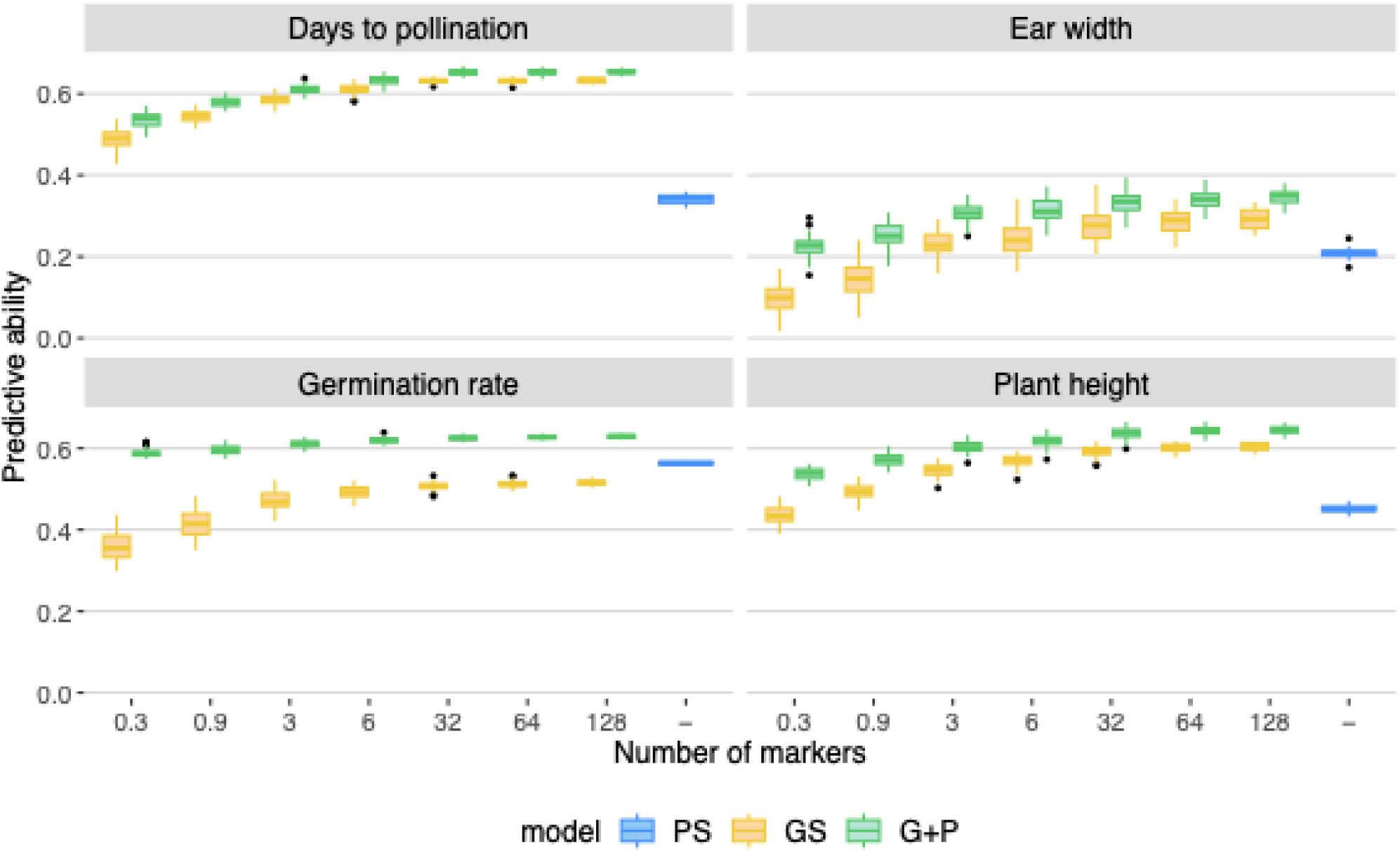
Predictive ability of three different models: genomic selection (GS), phenomic selection (PS), and a combination of both (G+P). The traits were evaluated in 2021. The x-axis shows the number of markers used to calculate the genomic relationship matrix for G and G+P approaches. To create the subsets of markers, six different set numbers (500, 1500, 5000, 10,000, 50,000, and 100,000) were randomly selected from an initial set of 200k. During the calculation of the relationship matrix, the initial set was filtered down to 128,202, and these subsets were represented on the x-axis as the average number of SNPs in thousands: 0.3, 0.9, 6, 32, and 64. For PS model, 730 wavelengths were used

### 3.4 Single gene starch mutation can affect accuracy

Given the observed separation between individuals with *su1* mutation and *sh2* mutation in the PCA based on skNIR data (Fig. 2), we explored prediction ability within these mutation groups. For *su1*, there were 364, 429, and 427 individuals in 2019, 2021, and 2022, respectively, whereas, for *sh2*, there were 150, 194, and 193 individuals for the same years.

The results of within-group prediction, compared to those using the entire diversity panel set, varied by trait, year, and model (Fig. 7). Overall, the average prediction ability of P-BLUP for all traits remained the same (around 0.24) for the group with the *su1* mutation, which had about 32% fewer individuals than the whole panel. However, it decreased to 0.05 for the group with the *sh2* mutation, which had about 70% fewer individuals than the entire panel. G-BLUP showed similar results, with the entire panel having an average prediction ability of 0.40, the *su1* group having 0.41, and the *sh2* group decreasing to 0.21.

**Fig. 7.**
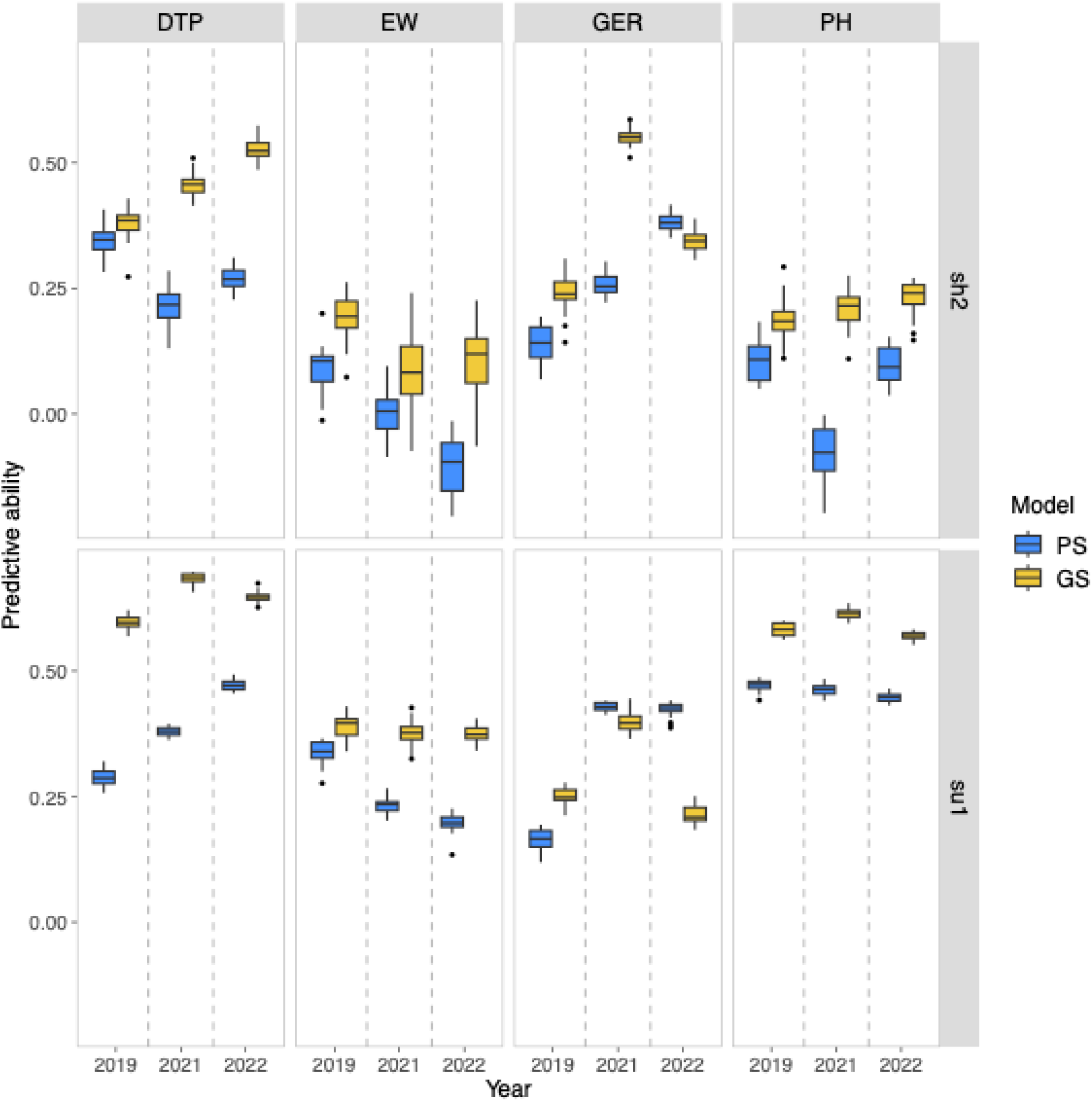
Predictive ability within groups in the diversity panel set with mutations in the *sugary1 (su1)* and *shrunken2 (sh2)* genes. Predictive ability is presented for genomic selection (GS), phenomic selection (PS), and a combined approach using both genomic and phenomic data (G+P) models. The traits evaluated over three years (2019, 2021, and 2022) were days to pollination (DTP), ear width (EW), germination (GER), and plant height (PH). Phenomic data comprised NIRS spectra averages from 2019 and 2020, processed using the Standard Normal Variate first derivative preprocessing method

For phenomic prediction, the most affected traits for the *su1* group compared to the whole panel were AC and SC, which decreased by around 0.20. Germination was also affected, with a decrease ranging from 0.06 in 2019 to 0.14 in 2022. Some traits, like DTP, increased, achieving 0.47 compared to 0.35 for the entire panel set. For individuals with the *sh2* mutation, some of the most affected traits were PH and AC, with a decrease in predictive ability of 0.4 and 0.35 compared to the whole panel, respectively.

### 3.5 Germination rate is not only explained by the starch composition

The phenomic selection model demonstrated a relatively high prediction ability for germination rate. We investigated the correlation between GER predictions and kernel composition traits within each group to determine if the model, while accurate for germination rates, may have some correlation with compositional traits of the kernel, such as starch or sugar content, and exhibit bias towards these components. Kernel composition was determined using a pre-existing PLS model with skNIR data for 237 individuals in the diversity panel, comprising 149 with *su1* mutation and 82 with *sh2* mutation

The correlation between germination rate and the kernel composition traits varied from −0.23 for glucose content to 0.29 for kernel weight, while correlations with starch content ranged from 0.18 to 0.24 (see Fig. 8 and S10). Overall, for the *su1* individuals, moving from BLUES of germination rates to predicted germination rates using PS increased positive correlation with total sugar, sucrose content, glucose content, starch content, phytoglycogen, and kernel weight. The only trait that did not show significant change was oil content. When using GS to predict values, there were no or small changes in correlation. For example, the correlation between BLUES of germination rates and glucose content is −0.19. The estimated value based on phenomic prediction is −0.15 and for genomic prediction is −0.08, while for starch content it is 0.23, 0.38, and 0.17 for BLUES, phenomic, and genomic estimation, respectively (Fig. 8). While for *sh2*, the pattern was not the same, there was an increase in positive correlation for kernel weight, oil content, starch content, sucrose content, and phytoglycogen, although small and less pronounced than for the *su1* individuals. There was none or a small increase in the negative correlation for glucose and total sugar content. The results vary slightly between years, but overall, they are similar (Fig. 8 shows the results for 2021, and Fig. S10 shows the results for the other years).

**Fig. 8.**
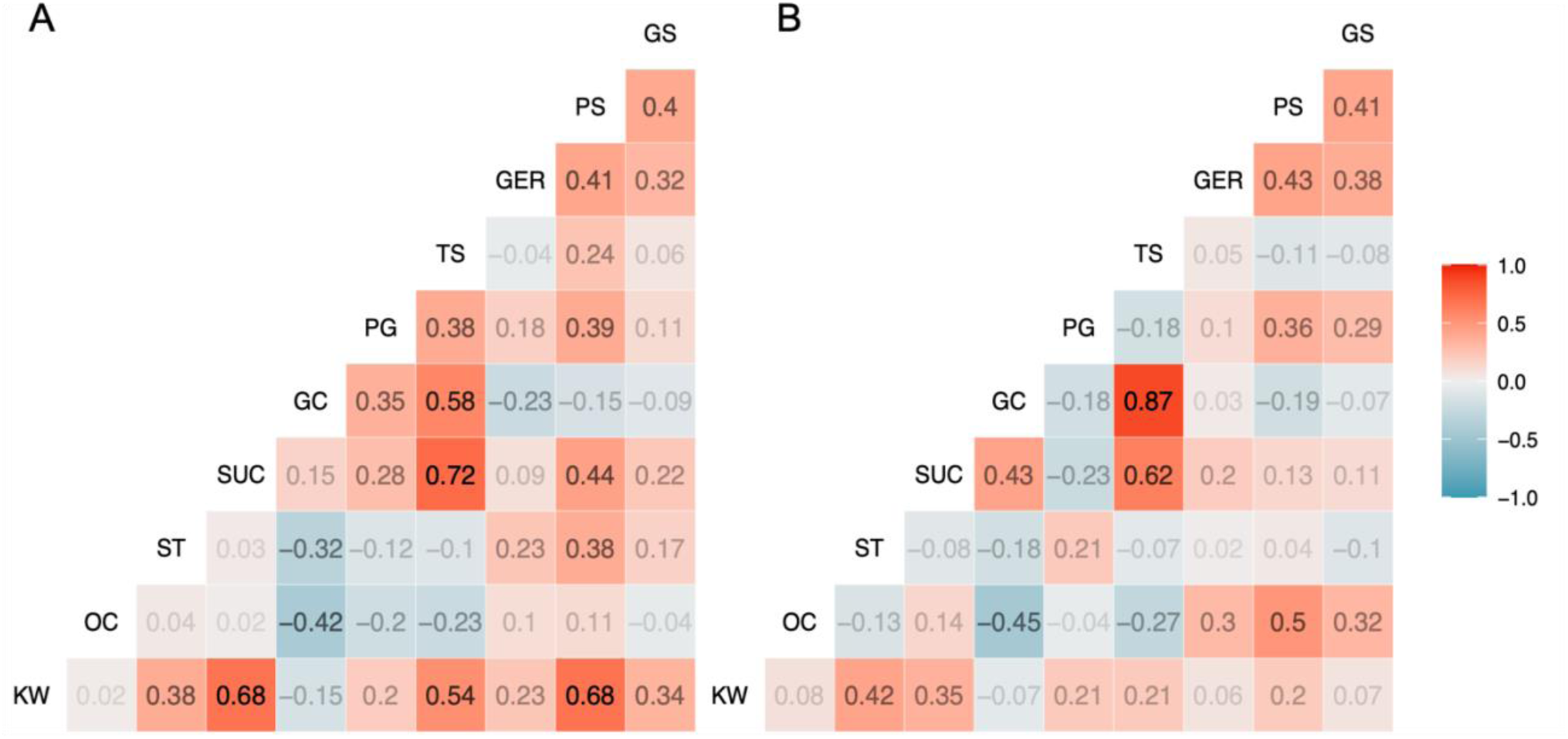
Correlation matrix displaying the relation between genomic and phenomic predictions with kernel composition traits. The kernel composition traits were measured using the NIRS with PLS method, which includes kernel weight (KW) (mg), oil content (OC,%), starch (ST,%), sucrose (SUC, mg), glucose (GC,%), phytoglycogen (PG, %), and total sugar (TS,mg). The GER represents the BLUEs of the germination rate observed in 2021. The PS and GS represents the predicted values for the germination rate obtained from phenomic and genomic predition (i.e. PBLUP and GBLUP), respectively. The predictions were made within groups in the diversity panel with mutations in the *sugary1* (A) and *shrunken2* (B) genes. Each cell in the matrix shows the Pearson correlation coefficient between pairs of values, with positive correlations displayed in red and negative correlations in blue. The PS model used the average NIRS data collected in 2019 and 2020 and the Standard Normal Variate first derivative preprocessing method

### 3.6 Phenomic prediction has the potential to predict germination rate before plating

We used the PS model trained with the diversity panel set to predict and select DH lines for germination before planting. The results showed that DH lines predicted to have high and low germination rates based on the PS model demonstrated high and low germination when planted in the field (Figs. 9A and 9B). The correlation between the observed and predicted values was high (0.84). The difference between the high and low groups was highly significant (p-value <0.05) and the correlation between the observed and predicted values was high (0.84). Additionally, we compared four individuals (two with low and two with high germination rates) whose seeds were produced in two locations (Chile and Idaho). We observed that seed source had an influence on germination rate, with seeds from Idaho performing better than those from Chile. However, the separation between the high and low germination groups was still evident even though predictions were generated only using the seeds from Chile (Fig. 9D).

**Fig. 9.**
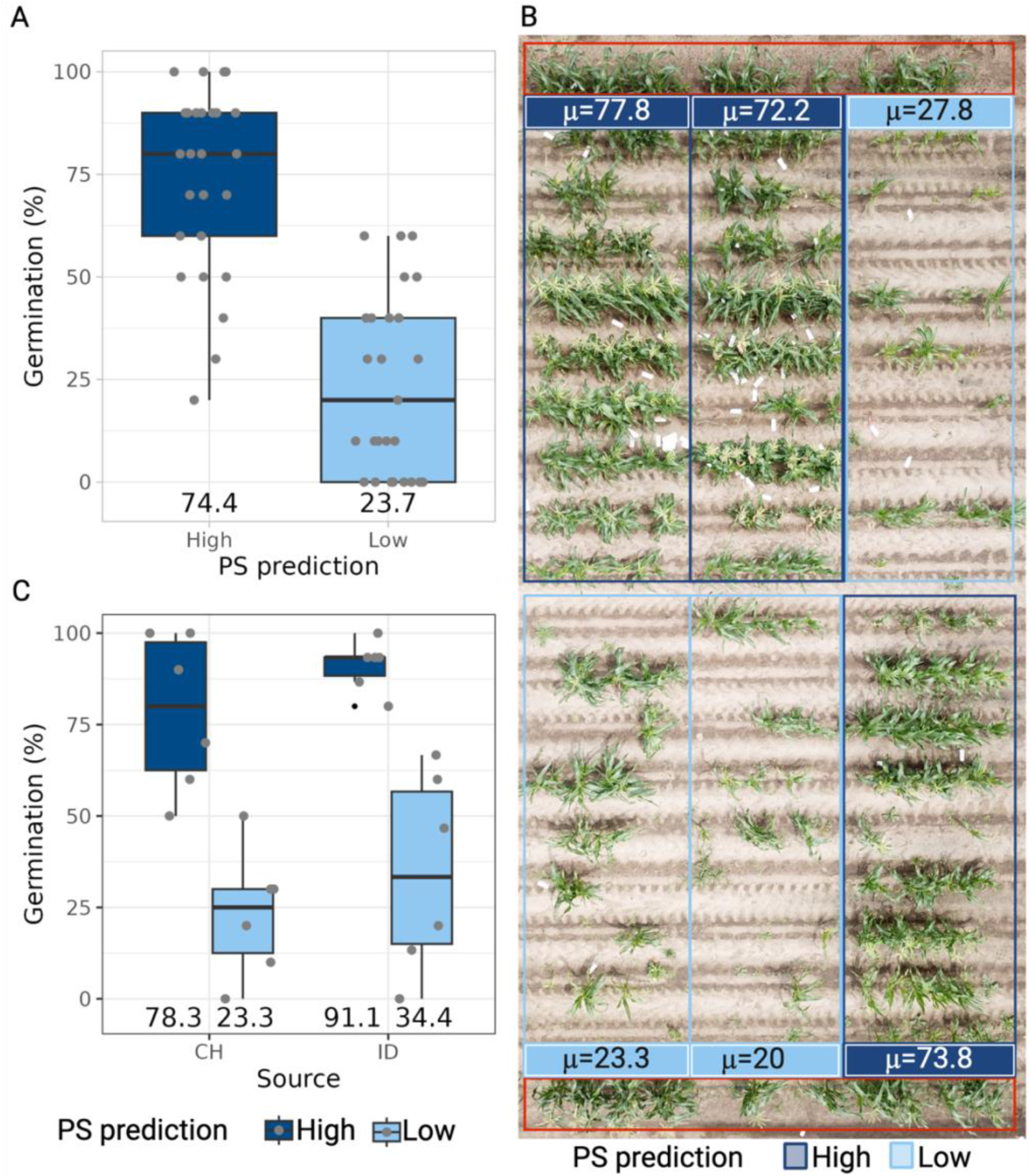
Results for predicting and selecting doubled haploids (DH) lines for germination trait before planting using PS model trained with the diversity panel set. Individuals with high and low germination rates were selected based on the predicted values. (A) presents the boxplot for the germination rate with the x-axis dividing individuals selected based on PS prediction, and the y-axis is the observed germination rate. Gray dots represent values per genotype. (B) displays a photo of the top of the field with a legend indicating if germination was predicted to be low or high based on PS, followed by the repetition mean. Inside the red shape are the borders. (C) presents the boxplot only for individuals with germination rates from seeds produced in Chile (CH) and Idaho (ID). The x-axis divides individuals on seed source; the y-axis is the observed germination rate. The predictions of the PS model used skNIR-sorter data from CH kernels

## 4 Discussion

According to the breeder’s equation, which quantifies genetic gain (Δ𝐺), Δ𝐺 = 𝑖𝑟𝜎_𝑎_⁄𝐿, where *i* represents the selection intensity, *r* is the selection accuracy, 𝜎_𝑎_ is the additive genetic standard deviation, and *L* is the cycle length (Desta and Ortiz, 2014). Genomic selection can enhance genetic gain by improving selection accuracy, increasing selection intensity, and shortening cycle length through early selection (Zhang et al. 2017; Ferrão et al. 2017; Xu et al. 2020; Beyene et al. 2021). Phenomic selection (PS) has emerged as a promising and cost-effective tool for predicting complex traits (Rincent et al., 2018). In addition to its potential to increase selection intensity and efficiency, some results indicate that PS could achieve selection accuracy similar to GS (Brault et al., 2022; Cuevas et al., 2019; Krause et al., 2019; Lane et al., 2020; Rincent et al., 2018; Robert et al., 2022a; Weiß et al., 2022; Zhu et al., 2022, 2021). To further improve genetic gain, it remains to be determined if PS can reduce cycle length and if early selections based on PS are feasible. To address this gap, our study utilized skNIR data to evaluate the predictive ability of PS in sweet corn.

We built a phenomic relationship matrix using a diverse panel with 24 traits observed over three years, including traits with varying heritability. Our results indicated that the predictive ability of PS was promising but trait dependent. While some traits could be predicted with relatively high accuracy from skNIR data, exceeding 0.35, genomic prediction generally outperformed phenomic prediction. Most PS studies have attempted to predict only a few traits, generally finding that phenomic prediction generates comparable or higher predictive ability than genomic prediction (Brault et al., 2022; Cuevas et al., 2019; Krause et al., 2019; Lane et al., 2020; Rincent et al., 2018; Robert et al., 2022a; Weiß et al., 2022; Zhu et al., 2022, 2021). The difference in PS prediction accuracy among different studies (including this one) could be related to the difference in plant tissues used to obtain phenomic data. Here, we used whole maize kernels. Intact whole kernels are relatively large and compositionally asymmetric, which could impact prediction accuracy compared to more uniform samples such as ground homogenized kernels (Desalvio et al., 2024). There may also be lower sensitivity of single kernel vs bulk kernel NIR (Hacisalihoglu and Armstrong 2023). However, the advantages, such as low cost, high speed, and non-destructive process, may still make it worthwhile to use individual, viable kernels despite lower prediction ability rather than grinding a kernel sample.

Our results indicated that vegetative traits were generally more accurately predicted than ear-related traits, a pattern seen in both PS and GS. Germination was one of the few traits for which phenomic prediction showed similar or higher accuracy compared to genomic prediction over all the years. Additionally, traits apparently unrelated to seed composition, such as plant height and days to pollination, exhibited relatively high prediction ability. The multitrait model did not improve prediction ability, which aligns with Peixoto et al. (2024) study on sweet corn hybrids. One advantage of the multitrait approach is leveraging the correlation between traits, but in our case, some traits had weak correlations, thereby limiting any potential advantage. The multi-trait approach may increase accuracy within specific trait groups, such as ear and vegetative traits, with stronger correlations.

### 4.1 Environment and seed source are not the main effects on phenomic prediction ability

Another important point to address is the proportion of environmental effects captured by the spectra. Since we used seeds in our study, we likely captured minimal environmental effects compared to, for example, scanning leaves in the field. This is reflected in the variance components along the spectrum, showing significant genetic variance and minimal GxE interaction, literature, also observed in other studies (Rincent et al. 2018; Zhu et al. 2021, 2022). However, we observed that the environment in which the kernels for the skNIR data were collected impacted the prediction ability values, although the differences were generally small (Fig. 3).

Furthermore, we noticed higher predictive ability for specific traits such as germination when the kernels scanned by the skNIR were from the same seed source as what was planted for phenotypic traits. This could be explained by the fact that if the seeds scanned and planted were from the same environment, an environmental effect on the seed could also influence the measured trait, leading to a non-genetic correlation between NIRS and the trait. This could particularly impact germination, as this trait is more influenced by the source of the seed (Wulff 1995; Xue et al. 2021). This may also explain why germination had a lower predictive ability in 2019 for GS and PS, as the set of seeds used to plant that year consisted of older seeds, affecting the germination potential.

To understand this better, we had scenarios where all the seeds planted for trait assessment came from a different year than the skNIR data used for prediction. This did not result in a clear pattern; even for germination, the prediction ability in this scenario was not always smaller (Fig. S8). Additionally, when validating the model with the DH seeds, values predicted using skNIR data from seeds sourced from Chile could accurately separate high and low germination groups for individuals planted with seeds sourced from Idaho, although this was performed with a small dataset. This finding indicates that minor variations in prediction ability, depending on the source of the NIRS data, could be due to spectra partially capturing environmental effects. However, the variation was too small to fully explain the good predictions, especially for traits less correlated with the sample, like plant height. This aligns with other studies’ findings that PS could be used to make predictions in independent environments (Posada et al. 2009; Robert et al. 2022b).

### 4.2 Potential impact of relatedness and endophenotypes on prediction accuracy

Rincent et al. (2018) highlighted that NIRS can capture polygenic relationships and tag major QTLs, which are related to factors behind GS accuracy, such as relatedness and linkage disequilibrium. As we mentioned in the introduction, this gives rise to two hypotheses: i) the predictive power of such an approach can come from the ability of a multivariate phenotype to estimate genetic relationships, which can, in turn, be used to predict any phenotype of interest; or ii) the ability to exploit endophenotypes by associating them with the target trait.

When comparing the PLS model to the P-BLUP approach, we found that P-BLUP generated similar or better results, indicating that NIRS could capture genetic relatedness, which led to better performance of phenomic selection. However, these differences were generally not high, suggesting that other factors may contribute to the observed performance. This aligns with studies that have indicated PS can make accurate predictions in unrelated individuals, while genomic prediction was affected by the lack of relatedness, suggesting that PS relies less on relatedness for prediction compared to GS (Weiß et al., 2022; Zhu et al., 2022, 2021).

For the second hypothesis, the ability to exploit endophenotypes by associating them with the target trait, NIR spectra could act as a proxy for the target trait through direct or indirect correlation (Robert et al., 2022). In the direct case, the target trait is related to the sample analyzed, such as when NIR spectra collected from the kernel are used to predict kernel composition, which aligns with the traditional application of NIRS. In the indirect case, the target trait is not directly linked to the sample but is correlated with its biochemical composition or characteristics. The multivariate estimate of these parameters with NIRS may be a more accurate synthesis of complex traits where the non-linear interaction underlies the prediction. This is particularly relevant for traits like germination, where NIR spectra can predict starch and sugar content, which can be correlated with germination. However, our results showed only a small correlation with kernel composition traits and no clear pattern for starch and sugar content, indicating that their direct correlation with germination would not be the only factor behind phenomic prediction ability.

We also hypothesize that PS could exploit endophenotypes by capturing genetic information from unrelated traits via linkage with the genes affecting the observed absorbance/reflectance at certain wavelengths. This could explain why traits not directly related to the kernel composition, such as days to pollination or plant height in the future grow out, can still yield relatively accurate predictions. This would align with results from Rincent et al. (2018) that indicate that NIR spectra can tag major QTLs.

The influence of each factor likely varies depending on the trait being predicted. Traits such as germination, which are more correlated with the sample may rely more on the association between the trait and the phenomic data. This also applies to traits like phosphorus concentration in the seed, which showed a high prediction ability for PS (Weiß et al., 2022). In contrast, less related traits such as plant height or ear width might benefit more from relatedness. Further research is needed to explore this in more detail. However, even in GS, it is not straightforward to determine how much each factor influences the accuracy (Habier et al. 2013).

Additionally, we found correlations between traits and wavelengths, with some overlap in trait correlation and predictive ability. This supports the second hypothesis, suggesting that when a trait is more closely related to the sample analyzed, NIRS data can better predict the target traits. Similarly, in bread wheat, Kruse et al. (2019) performed PS with hyperspectral time-point-derived relationship matrices to predict grain yield and observed that prediction accuracy was highly correlated with the strength of the relationship between hyperspectral reflectance and the trait. Considering the hypothesis of associating target trait with endophenotypes, whether from linkage or as a proxy, one potential method to investigate this association is to examine the relationship between higher correlations and improved prediction accuracy. This can help to clarify the genetic improvement achieved when using PS by using it as an indirect selection tool. By considering the NIR spectra as an index (I), we can formalize this concept as follows:

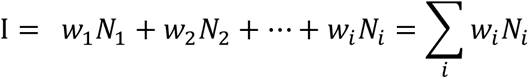

where *w_i_* would be the weight of the wavelength *i* and the N would be the absorbance of the wavelength *i*, what can be obtained with ridge regression best linear unbiased prediction model. Then, according to Falconer and Mackay (1996) the correlated response of a trait *Y* based on the NIR spectra as an index (I) would be as follow:

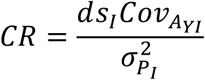

where 𝑑𝑠_𝐼_ is the selection differential for the index, 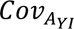 is the additive genetic between the index and the trait y and 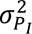 would be the phenotypic variance of the index. As I is the NIR spectra, the covariance and variance terms become 𝐶𝑜𝑣(𝑌, ∑_𝑖_ 𝑤_𝑖_ 𝑁_𝑖_) and 𝑉𝑎𝑟(∑_𝑖_ 𝑤_𝑖_ 𝑁_𝑖_), respectively.

### 4.3 Phenomic prediction within the two important mutation groups for sweet corn production

In the PCA analysis, we observed that the first two principal components (PCs) of the NIR spectra from the diversity panel separated the population into two groups related to the mutations in the *su1* and *sh2* genes. Although the differences occur at a single locus, which would likely not be captured by PS (Zhu et al., 2022), they affect seed composition. Therefore, this major QTL is likely related to a specific signal in the NIR spectrum.

When PS models were constructed within each group, the prediction results showed that the specific genetic mutation affects trait prediction differently. Since germination is partly associated with kernel composition traits, the decrease in accuracy when predictions were made within groups indicates that within-group population structure was confounding predictive ability. However, reduced accuracy could also be related to reduced sample size and, consequently, a smaller training set within groups (Zhu et al., 2022). For instance, predictions for germination were validated in DH lines that only contained the *sh2* group and still generated good results.

### 4.4 Enhancing germination rate through phenomic prediction in sweet corn

In sweet corn, germination rate is crucial because breeding for increased sweetness can lead to problems in germination and emergence due to mutations in endosperm starch synthesis (Lertrat and Pulam 2007; Revilla et al. 2021). This raises the question of whether prediction ability is influenced by kernel composition and whether selection for high germination rate with PS could unintentionally select for less sugar. For instance, a study on wheat found that although the PS model generated good results for predicting grain yield, it could lead to unintentional selection for lower protein content due to the negative correlation between yield protein content (Dallinger et al. 2023). This can be a limitation of PS if prediction selects undesirable traits.

Considering the division of the diversity panel into two groups of starch biosynthesis mutations, *su1* and *sh2*, if the selection is based on values obtained by the PS model, our results showed an increase in correlation with some compositional traits, but this was not straightforward. There was usually a positive correlation with starch content but no consistent increase in negative correlation with sugar components. This suggests that while predicting using skNIR spectra may not lead to selecting genotypes with less sugar content, it can introduce some bias toward kernel compositional traits. This indicates that the PS model could be used in this case, but the potential for undesired selection should be carefully considered.

Given the promising results of PS for germination rate in the diversity panel and the importance of this trait for sweet corn breeding, we performed a validation with a new dataset to assess the feasibility of this approach. We tested the model calibrated with the diversity panel by predicting germination rates in a new set of DH elite genotypes, all carrying the *sh2* gene. These previously untested lines had kernels scanned by the skNIR-sorter to obtain their phenomic breeding values. Based on these values, lines with low and high germination rates were selected and planted. The results confirmed that these individuals exhibited the expected opposite germination values, validating the use of PS for germination rate in sweet corn breeding. Since selection for increased sugar content may lead to poorer germination, a method to scan and remove those lines with low germination prior to planting would benefit the program by saving resources.

### 4.5 How to Integrate Phenomic Selection into the Breeding Pipeline?

The use of phenomic selection (PS) will depend on the crop and the trait analyzed. Some studies have shown high prediction ability for PS, but this was not true for all traits we analyzed. If PS exhibits high prediction ability, it could be used alone to rank individuals and assist breeders in the selection process. In cases where GS models perform better, it becomes a trade-off between intensity and accuracy.

The results of this paper indicate that skNIR spectroscopy could be used to scan seeds, perform prediction, and then plant. Our results with DH lines indicated that this application showed promising results in identifying low-germination individuals who would likely not advance in field trials. As pointed out by Rincent et al. (2018), PS models can be used to eliminate a proportion of selection candidates, even if prediction accuracy is low. The predictive ability of PS would determine the proportion of individuals that can be confidently filtered out without losing the best candidates. This application is particularly promising given the low-cost NIRS acquisition, often routinely conducted in some crops, thus limiting the number of individuals before planting and before more expensive screenings. For example, if a GS model is used to complement the PS model, this approach can save resources by reducing the number of individuals needing genotyping.

Another possible application would be to use PS combined with GS (G+P model) to increase the accuracy of selection. In this study, we used a multi-matrix model, including phenomic and genomic-based relationship matrices as separate terms. This model generated similar or better results than using skNIR or markers alone. This indicates that genomic and phenomic datasets captured different features, which can also be observed in the low correlation between the relationship matrices. Zhu et al. (2021) also found a low correlation between NIRS and marker matrices, and the G+P model showed better predictive ability, aligning with our findings and those of Krause et al. (2019) and Robert et al. (2022).

The combined model (G+P) increase relative to GS alone was particularly higher when low marker densities were used. The genomic selection based on a large number of markers may not be practical for routine application, especially for small breeding programs. We explored performing the predictions with different numbers of markers, and the results indicate that prediction ability generally shows a considerable increase up to 10,000 markers. In cases where PS alone has low accuracy, combining P+G could lead to high accuracies using fewer markers, as we observed high accuracies with just 500 markers. Computational and imputation methods have been developed to address low-density marker scenarios (Li et al. 2009). However, since NIRS is relatively easy to implement and is already routinely used in some breeding programs, it could be an effective way to increase the accuracy of genomic selection. We are dealing with a diversity panel; in the case of elite individuals, since there is less diversity, the decrease in accuracy with the reduction of the number of markers can be less pronounced.

Considering these two possible applications and breeding schemes involving inbred lines such as sweet corn, Fig. 10 illustrates these potential applications of PS using whole kernel NIRS spectroscopy based on our results. A first step in the program would be to scan all inbred lines generated from the F1 and perform phenomic selection to eliminate individuals that would likely not advance in the field trials (e.g., with low germination rates) before more rigorous or expensive screening. A second step would be to use PS combined with GS (G+P model) to improve selection accuracy. If the PS model were used to filter out the candidates, all genotyped individuals would have NIRS data available, leading to a straightforward implementation of the G+P model, which would allow for the use of fewer markers without additional steps.

**Fig. 10.**
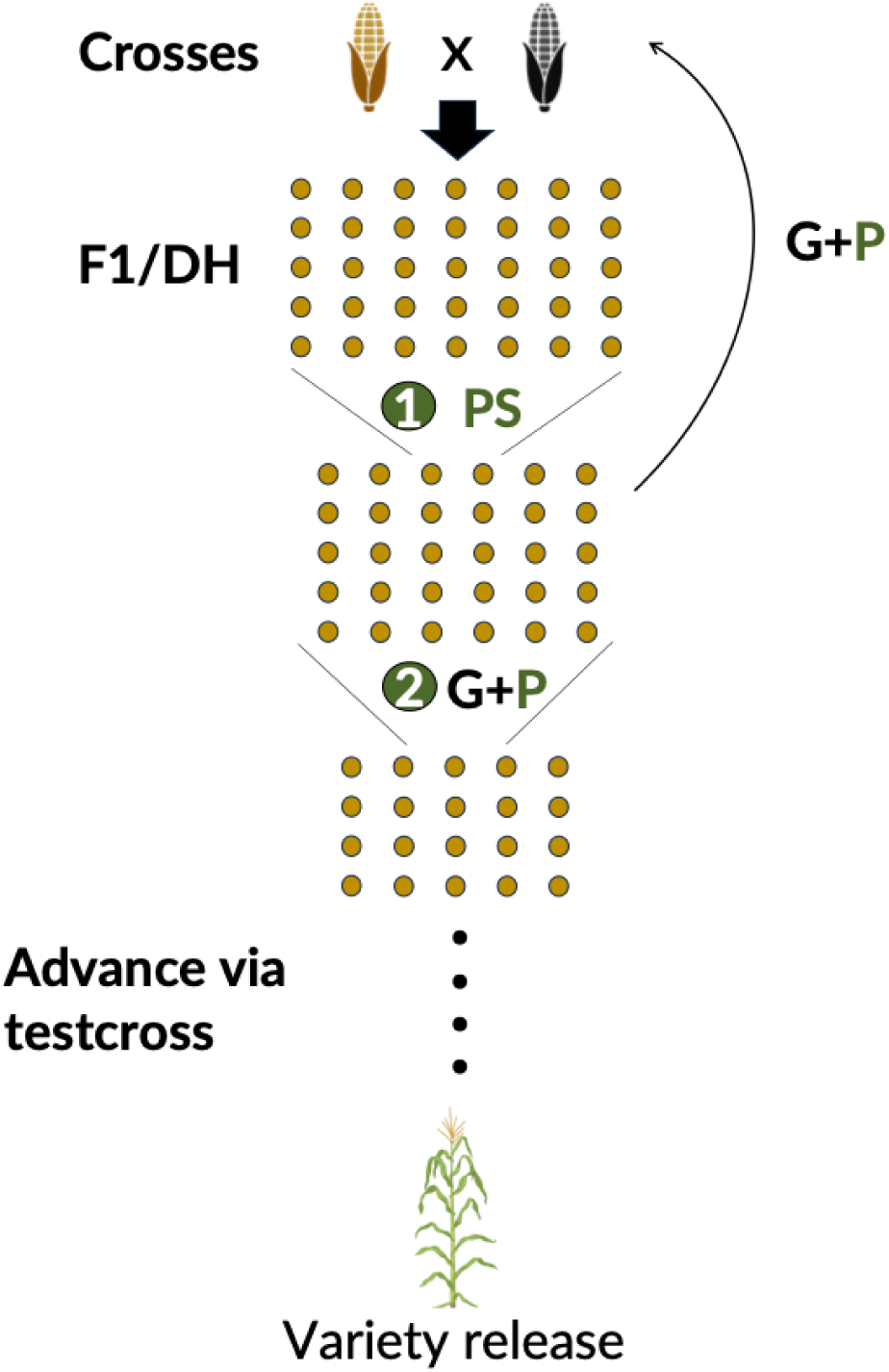
Schematic representation of a corn hybrid breeding pipeline with the potential application of phenomic selection (PS) from single-kernel NIR spectroscopy. Elite lines, represented by the corn ears, are crossed to produce F1 families, creating doubled haploid (DH) lines. In the first step, PS is used to screen and select the DH lines (represented by the yellow dots), eliminating a proportion of individuals. In the second step, the remaining individuals are genotyped. With both genomic and phenomic data available, a combined model (G+P) is used to select candidates that will advance to field trials via testcross, as well as the lines that will become parents of the next generation. Created with BioRender.com

An unexplored application in this paper is using the PS model in hybrids. If the NIRS acquisition is made directly in the single kernels, while this would be very useful, maternal effects could impact accuracy when scanning seeds before planting (Wulff, 1995). However, as discussed by Desalvio et al. (2024), it could also be valuable to predict the testcross performance of hybrids using skNIR spectroscopy data from the inbred lines (Albrecht et al. 2011; Peixoto et al. 2024b). It is also essential to consider non-additive and non-genetic effects when using PS. In this study, since we analyzed lines and obtained data from kernels, we do not have dominance effects affecting the predictive ability of PS, as well as minor non-genetic effects, as discussed earlier. However, further analyses are needed to better understand the effects of this method. Depending on the stage of the breeding process, these effects could compromise the estimation of breeding values. In some cases, such as testcross prediction in hybrid selection, capturing dominance could help better predict specific combining ability, and capturing environmental effects associated with genomic by environment interaction could benefit variety release.

## 5 Conclusion

While GS has been well-explored over the past 20 years, PS is still relatively new. The results indicate that the usefulness of PS in trait prediction for sweet corn depends on the specific trait. Although GS generally performed better than PS, combining both datasets produced the best results, especially with low marker density. The source of NIRS data had little influence on the phenomic prediction ability, but further studies are necessary to explore the nature of its accuracy. In addition, the validation with elite lines showed that obtaining data from whole seeds and performing selection before planting is feasible. Our results suggest that applying PS before or alongside GS for the sweet corn breeding program has the potential to accelerate genetic gains. Overall, PS is a promising tool, and this study gives insights into how best to leverage PS for optimal advantage.

## Acknowledgments

This work was supported by the National Institute of Food and Agriculture SCRI 2018-51181-28419 to AMS and MR, and AFRI 2019-05410, and USDA-NIFA 2022-51181-38333 to MR). Mention of trade names or commercial products in this publication is solely for the purpose of providing specific information and does not imply recommendation or endorsement by the U.S. Department of Agriculture. USDA is an equal opportunity provider and employer.

## Author contribution statement

RPG: Writing – original draft, Formal analysis, Methodology, Visualization.

MP: Writing – review & editing, Data curation, Methodology.

KL: Data curation, Investigation, Validation.

NS: Data curation.

JG: Data curation, Writing – review & editing, Resources.

PRA: Resources.

AMS: Funding acquisition, Writing – review & editing, Resources.

MR: Funding acquisition, Methodology, Project administration, Investigation, Resources, Supervision, Writing – review & editing.

