## Supplementary material 1 for "Integrating Phenomic Selection Using Single-Kernel Near-Infrared Spectroscopy and Genomic Selection for Corn Breeding Improvement"

**Journal: Theoretical and Applied Genetics**

**Supplementary Material 1**

**Table S1**. List of phenotypic traits evaluated in the diversity panel, with corresponding abbreviations and summary statistics for each year. The phenotypic mean is denoted by µ, and broad sense heritability is denoted by H^2^.

| Traits | Abbreviation | 2019 |  | 2021 |  | 2022 |  |
| --- | --- | --- | --- | --- | --- | --- | --- |
|  |  | µ | H^2^ | µ | H^2^ | µ | H^2^ |
| Days to pollination | DTP | 58.95 | 0.62 | 62.11 | 0.82 | 60.31 | 0.69 |
| Ear height | EH | 32.65 | 0.75 | 29.3 | 0.66 | 26.04 | 0.60 |
| Plant height | PH | 140.33 | 0.82 | 127.29 | 0.76 | 119.27 | 0.78 |
| Flag leaf height | FLH | 106.48 | 0.8 | 95.49 | 0.59 | 87.64 | 0.78 |
| Tassel extension | TE | 33.48 | 0.63 | 31.92 | 0.18 | 31.62 | 0.35 |
| Tiller number | TN | 1.39 | 0.5 | 0.71 | 0.47 | 0.86 | 0.38 |
| Leaf angle | LA | 44.75 | 0.65 | 47.83 | 0.69 | 48.9 | 0.44 |
| Ear length | EL | 127.09 | 0.71 | 12.99 | 0.69 | 12.08 | 0.57 |
| Ear width | EW | 34.51 | 0.65 | 3.4 | 0.67 | 3.15 | 0.44 |
| Kernel row number | KRN | 13.16 | 0.74 | - | - | - | - |
| Germination | GER | 57.39 | 0.69 | 50.17 | 0.67 | 53.17 | 0.40 |
| Convexity | COV | - | - | 0.83 | 0.42 | 0.82 | 0.19 |
| Solidity | SOL | - | - | 0.94 | 0.39 | 0.93 | 0.24 |
| Taper | TP | - | - | 85.76 | 0.5 | 72.41 | 0.41 |
| Curvature | CUR | - | - | 15.58 | 0.21 | 14.05 | 0.13 |
| Tip fill | TPF | - | - | 0.91 | 0.3 | 0.93 | 0.26 |
| Kernel row fill | KRF | - | - | 0.13 | 0.28 | 0.11 | 0.25 |
| Tassel branch number | TBN | - | - | 8.95 | 0.59 | 9.05 | 0.64 |
| Anther color | AC | - | - | 2.34 | 0.77 | 2.55 | 0.63 |
| Days to silking | DTS | - | - | 62.57 | 0.8 | 61.74 | 0.70 |
| Silk color | SC | - | - | 2.47 | 0.85 | 2.58 | 0.72 |
| Shank length | SL | - | - | - | - | 7.6 | 0.47 |
| Number of ear shoots | NES | - | - | - | - | 1.65 | 0.24 |
| Husk appearance | HAP | - | - | - | - | 2.02 | 0.46 |

**Table S2. Doubled haploids predicted values of germination rate for selected individuals based on phenomic prediction.** The rank reflects their ranking among all 522 individuals scanned and predicted using the PS model. The $\hat{y}$ denotes the predicted value for the germination rate. The group represents high germination rates (above 60%) and low germination rates (below 60%).

| Selected individual | $\hat{y}$ | Group |
| --- | --- | --- |
| 41 | 78.86 | high |
| 46 | 77.82 | high |
| 53 | 76.16 | high |
| 61 | 75.29 | high |
| 63 | 74.79 | high |
| 64 | 74.69 | high |
| 70 | 73.88 | high |
| 79 | 73.37 | high |
| 457 | 54.72 | low |
| 481 | 51.74 | low |
| 487 | 50.02 | low |
| 488 | 49.94 | low |
| 493 | 49.31 | low |
| 496 | 48.64 | low |
| 498 | 48.26 | low |
| 508 | 43.63 | low |
| 511 | 42.00 | low |

**Table S3.** Number of individuals included in each analysis, after restricting the dataset to genotypes present in both the genomic and spectra data.

| Phenotypic data year | skNIRS data source | Number of individuals |
| --- | --- | --- |
| 2019 | 2019 | 489 |
|  | 2020 | 513 |
|  | Average from both years | 521 |
| 2021 | 2019 | 555 |
|  | 2020 | 631 |
|  | Average from both years | 642 |
| 2022 | 2019 | 552 |
|  | 2020 | 628 |
|  | Average from both years | 639 |

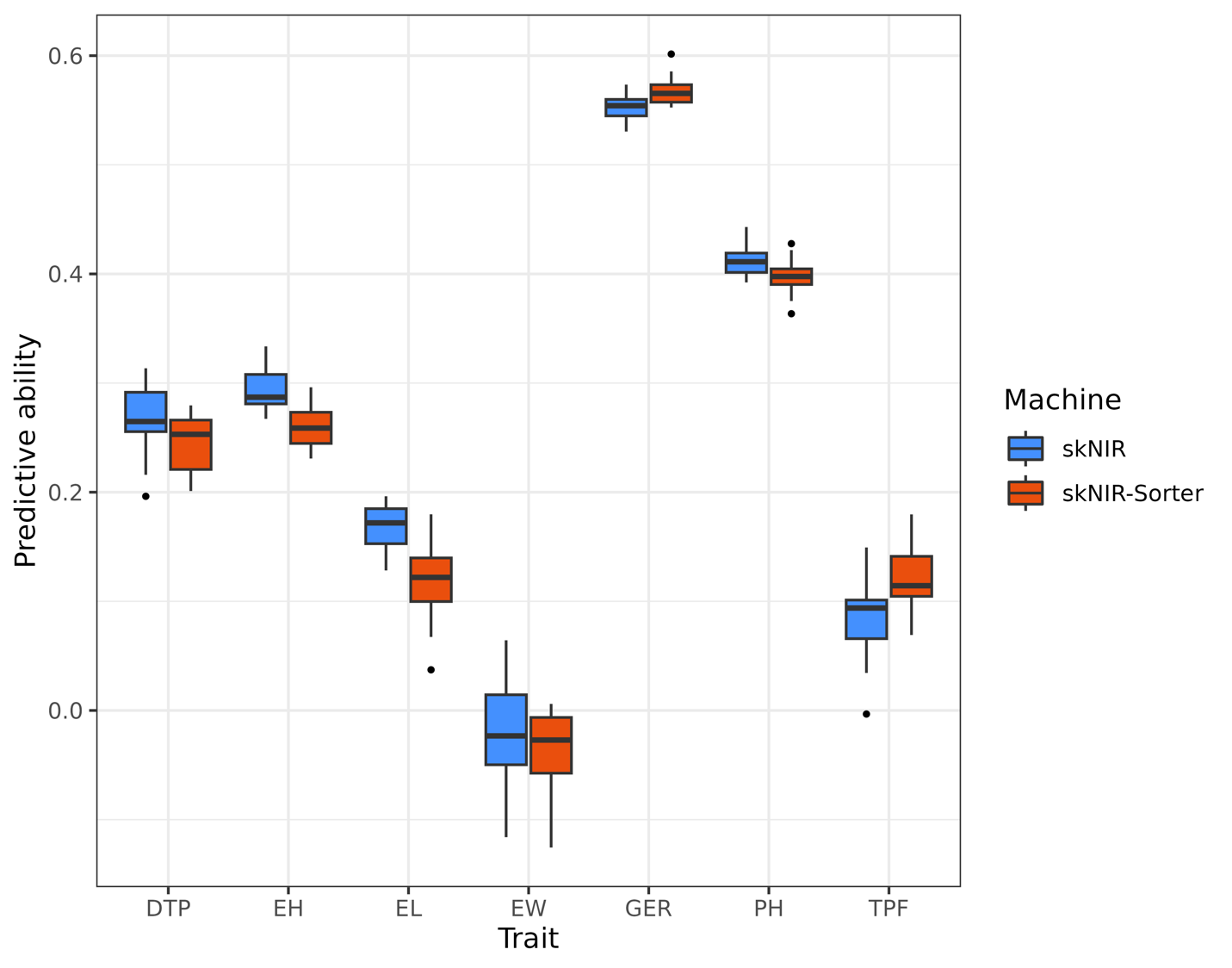

**Figure S1.** Boxplot of predictive ability of a model trained with NIRS data obtained from two NIRS machines: the single kernel near infrared (skNIR) and the single kernel near infrared sorter (skNIR-Sorter). The analysis involved 256 individuals from the diversity panel scanned by both devices. The dataset used for the NIRS for both devices came from the seeds harvested in 2020. The results were derived from 20 repetitions of fivefold cross-validation and are presented for phenotypic data assessed in 2022, including days to pollination (DTP), ear height (EH), ear length (EL), ear width (EW), germination (GER), plant height (PH), and tip fill (TPF).

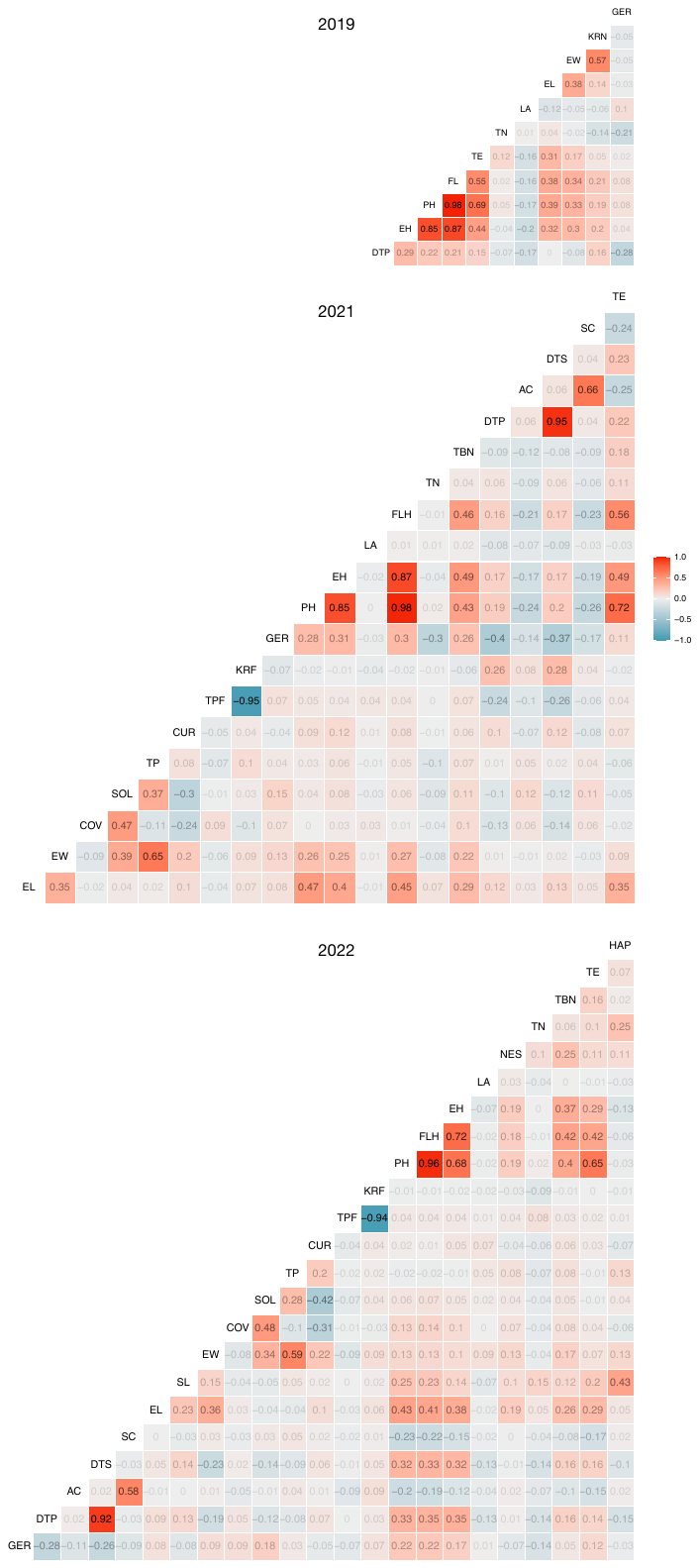

Figure S2. Phenotypic correlation matrix for traits in the diversity panel across 2019, 2021, and 2022. Each cell displays the Pearson correlation coefficient between pairs of traits, with color coding to denote positive (red) and negative (blue) correlation.

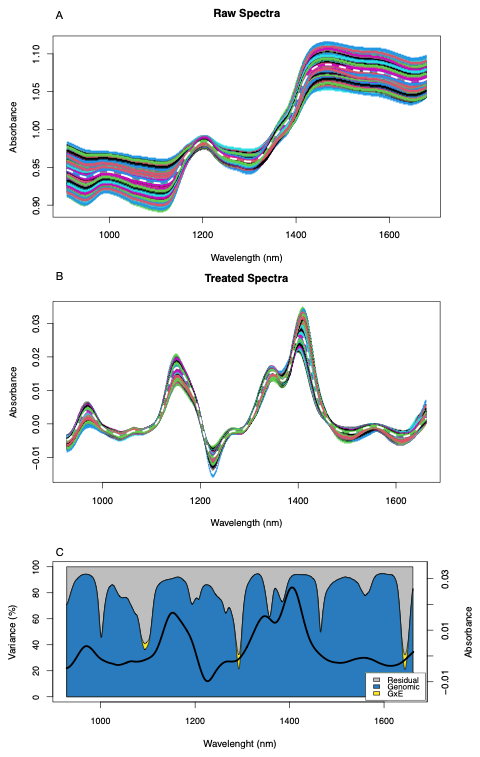

Figure S3. The (A) displays the raw NIRS data averaged from scans of the diversity panel in 2019 and 2020, while (B) shows the same data processed using the Standard Normal Variate first derivative approach. C) displays the proportion of genetic (blue), genetic by environment (yellow), and residual (grey) variances along the NIR spectrum. The standard normal variate first derivative average spectra is indicated in black.

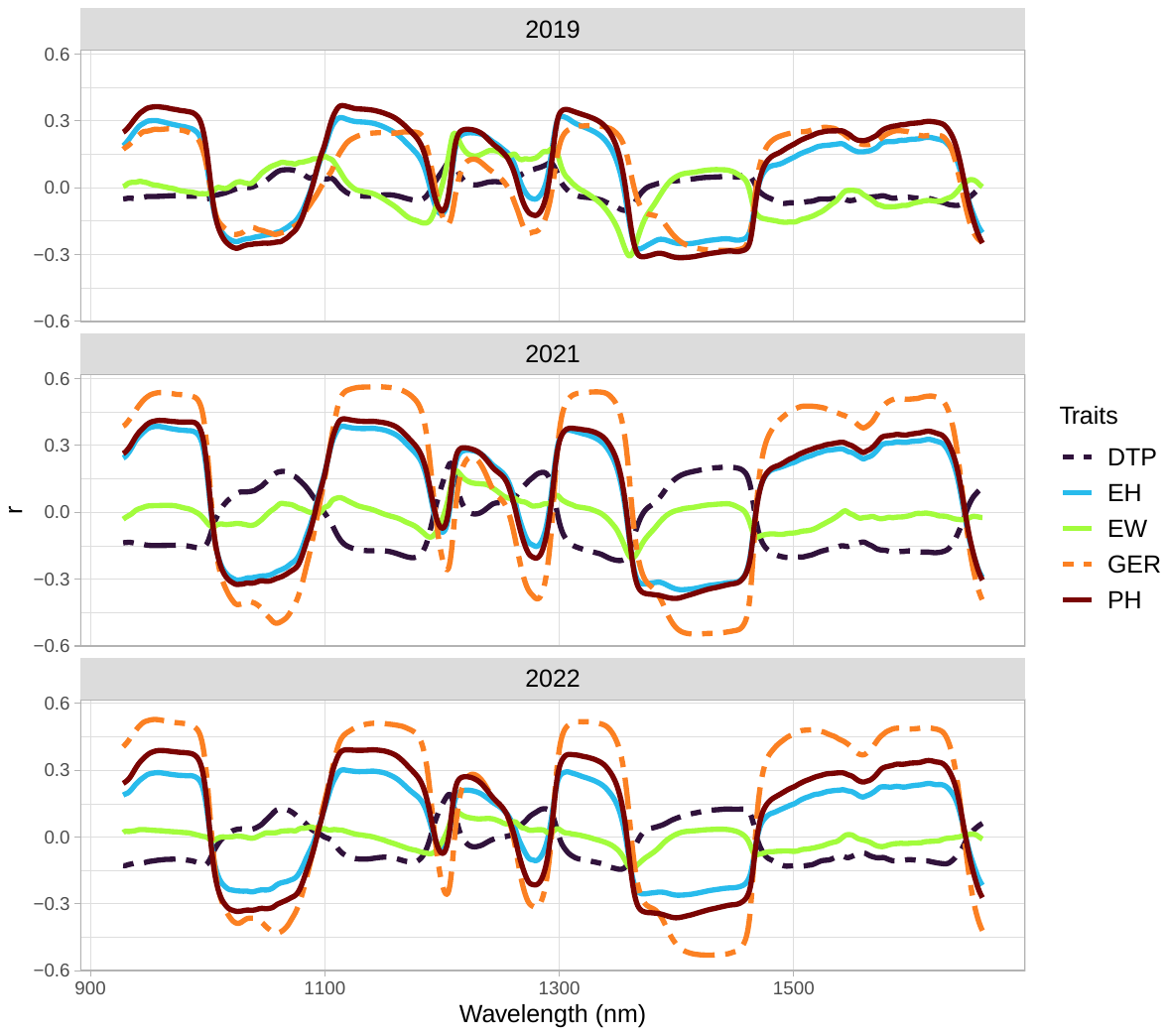

**Figure S4.** Correlation (r) between each wavelength and vegetative traits for the diversity panel in 2019, 2021, and 2022, respectively. Phenomic data comprised NIRS spectra averages from 2019 and 2020, processed using the Standard Normal Variate first derivative preprocessing method. The traits evaluated over three years (2019, 2021, and 2022) were days to pollination (DTP), ear height (EH), ear width (EW), germination (GER) and plant height (PH).

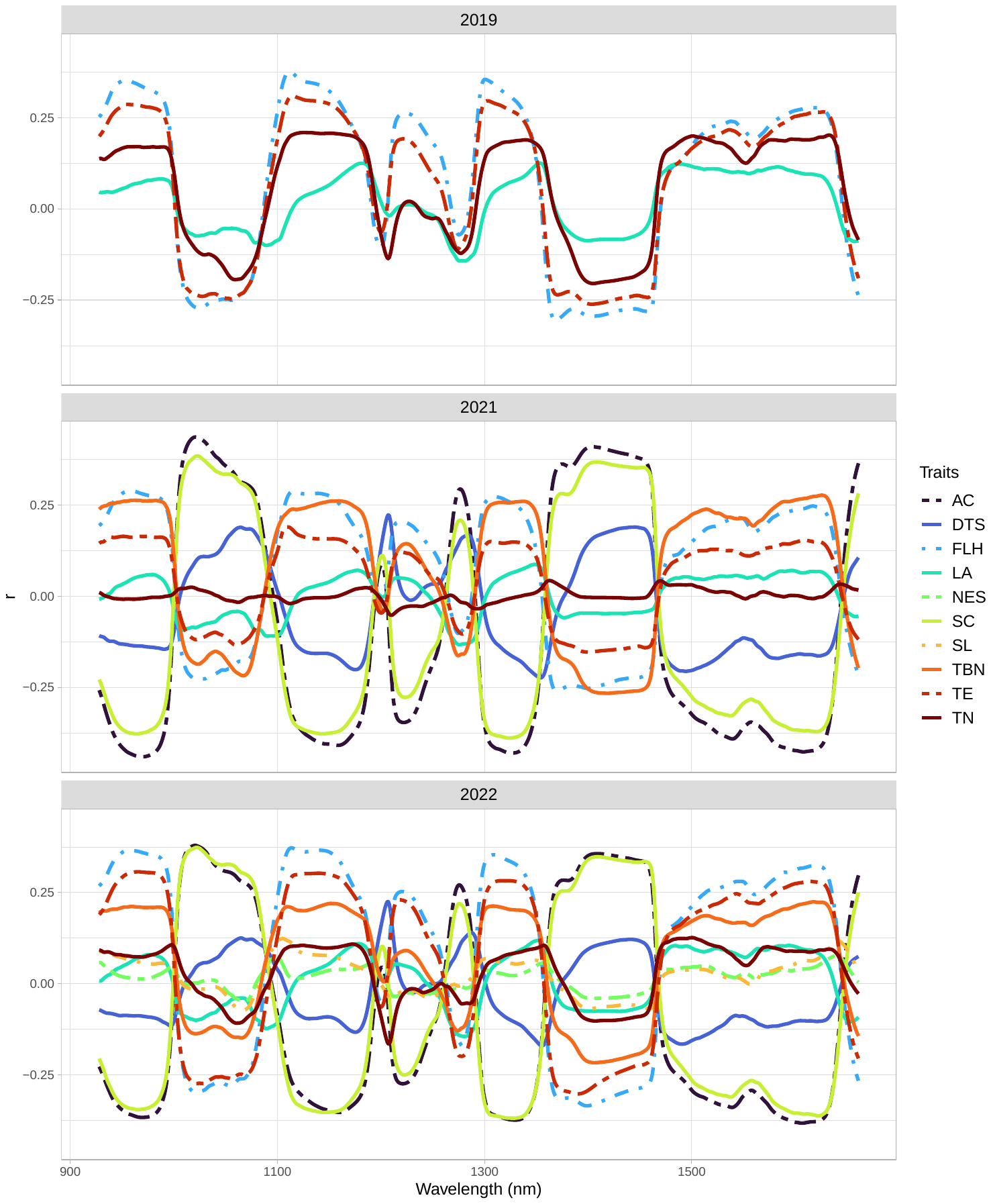

**Figure S5.** Correlation (r) between each wavelength and vegetative traits for the diversity panel in 2019, 2021, and 2022, respectively. Phenomic data comprised NIRS spectra averages from 2019 and 2020, processed using the Standard Normal Variate first derivative preprocessing method. The traits evaluated over three years (2019, 2021, and 2022) were: DTS: days to silking; FLH: flag leaf height; TE: tassel extension TN: tiller number; LA: leaf angle; TBN: tassel branch number; SL: shank length; AC: anther color; SC: silk color; NES: number of ear shoots).

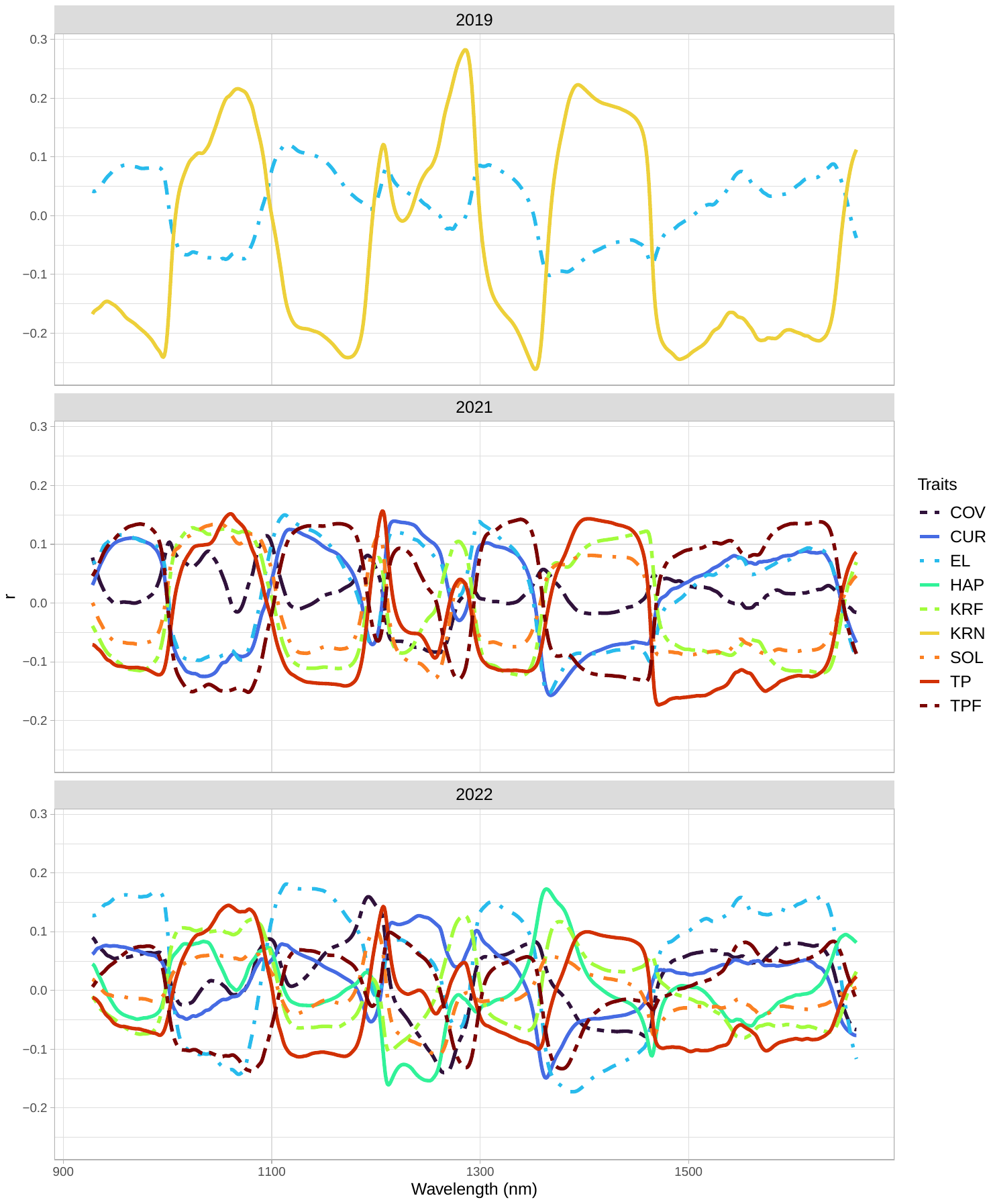

**Figure S6**. Correlation (r) between each wavelength and vegetative traits for the diversity panel in 2019, 2021, and 2022, respectively. Phenomic data comprised NIRS spectra averages from 2019 and 2020, processed using the Standard Normal Variate first derivative preprocessing method. The traits evaluated over three years (2019, 2021, and 2022) were: EL: ear length; TPF: tip fill; HAP: husk appearance; KRN: kernel row number; SOL: solidity; TP: taper; CUR: curvature; COV: convexity; KRF: kernel row fill).

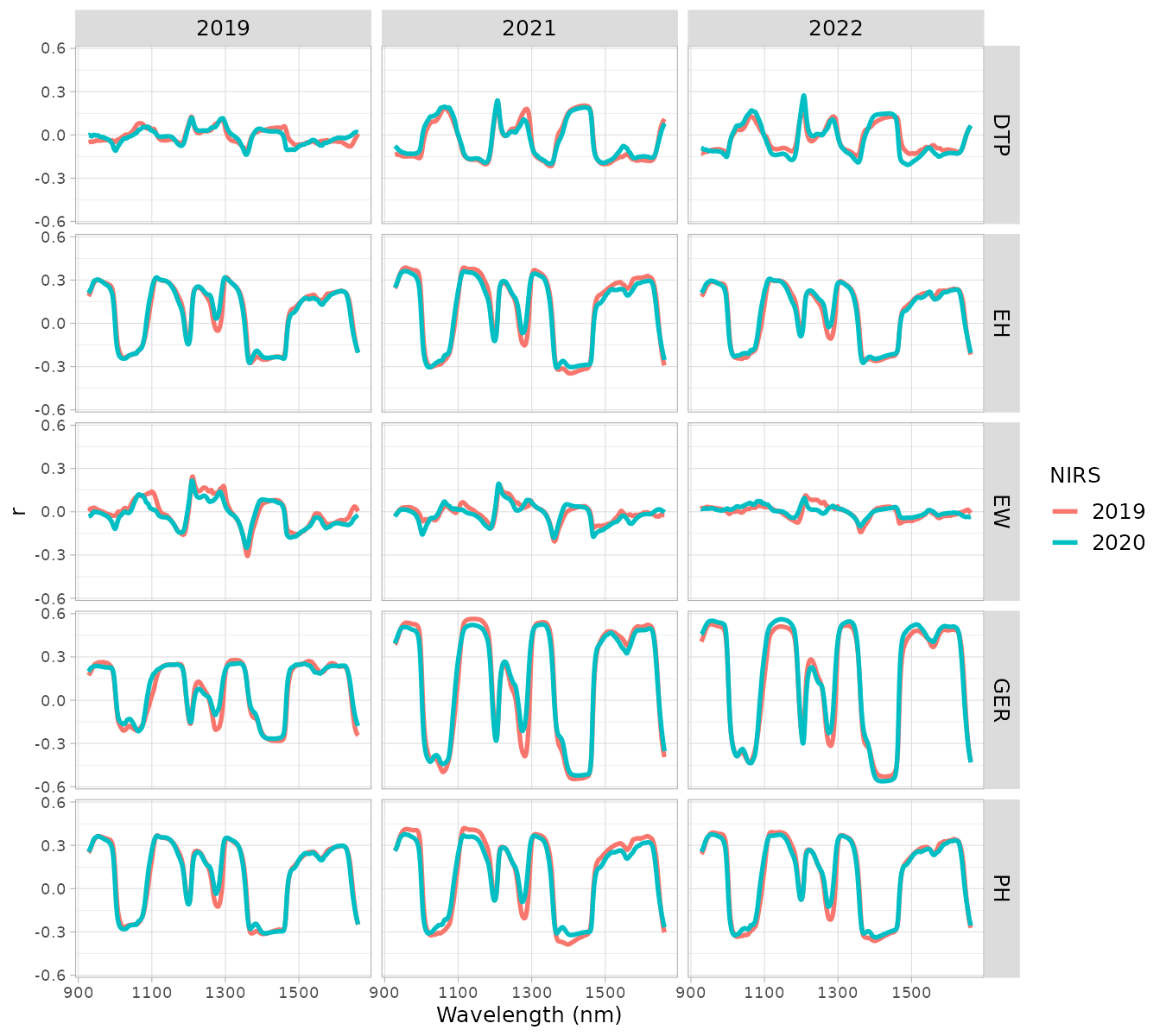

**Figure S7**. Correlation (r) between each wavelength and vegetative traits for the diversity panel using NIRS data from 2019 and 2020. in 2019, 2021, and 2022. Phenomic data was processed using the Standard Normal Variate first derivative preprocessing method. The traits evaluated over three years (2019, 2021, and 2022) were days to pollination (DTP), ear height (EH), ear width (EW), germination (GER) and plant height (PH).

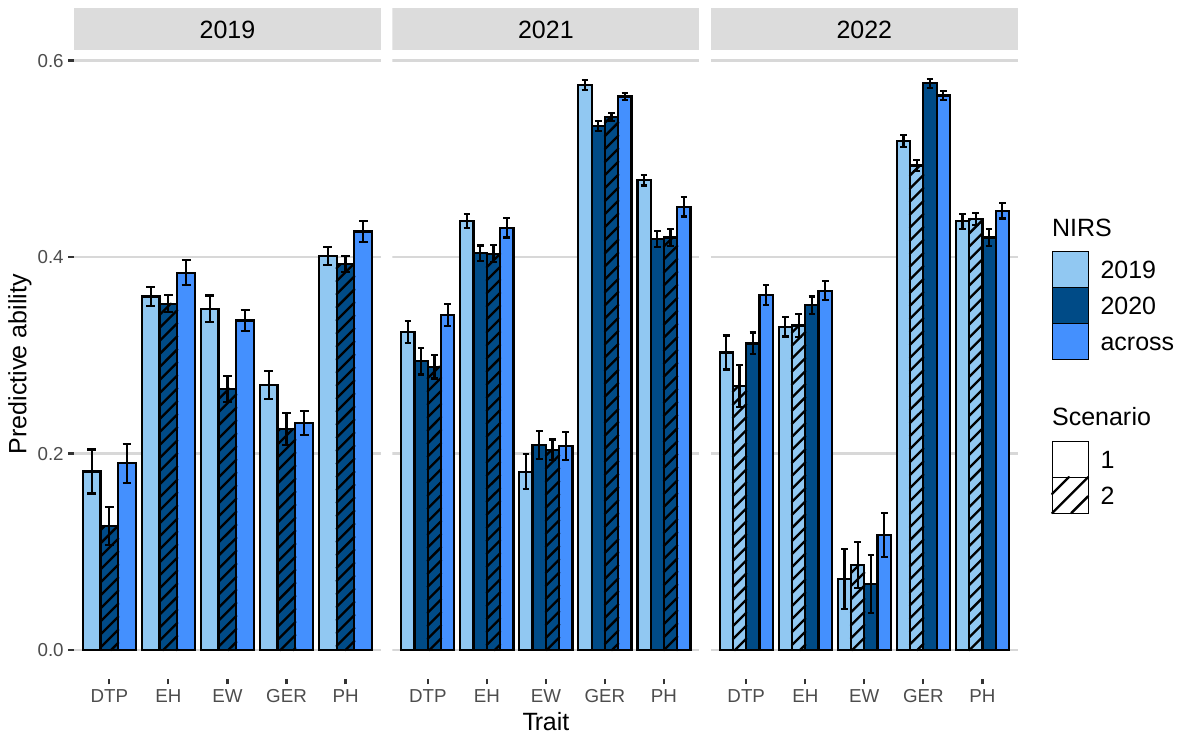

**Fig. S8** Predictive ability of phenomic selection (PS) using NIRS data collected in different years, 2019, 2020, and an average across the two years (across). Scenario 1 includes all available data sets. To account for potential environmental variation captured by skNIR data, we considered scenario 2, ensuring there was no overlap in seed source used for skNIR measurement and planted for phenotypic data in the analysis. The aim was to remove the environmental effect in the predictions due to the non-genetic correlation between the skNIR data and the phenotypic data. This means that the environmental conditions affecting the scanned kernels did not influence the traits measured in the plants. For predictions regarding phenotypic data in 2019, skNIR data from the 2020 kernel source was used with no additional steps, as no seeds from 2020 were planted for phenotypic evaluation in 2019. However, for 2021 and 2022, individuals were filtered to ensure different seed sources. The traits evaluated over three years (2019, 2021, and 2022) were days to pollination (DTP), ear height (EH), ear width (EW), germination (GER), and plant height (PH). Error bars are standard errors of means from 20 repetitions.

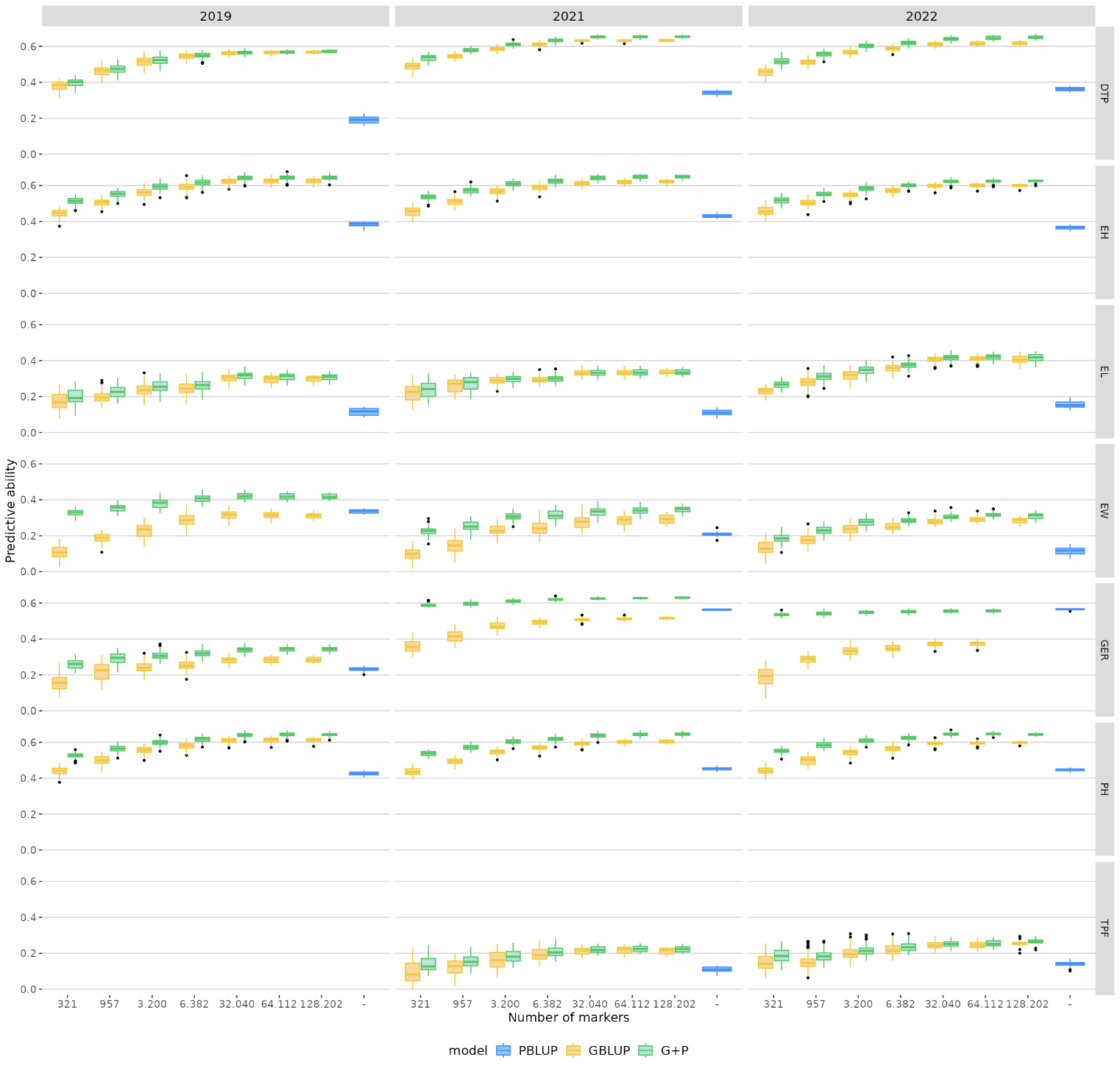

**Fig. S9** Predictive ability of three different models: genomic selection (GS), phenomic selection (PS), and a combination of both (G+P). The x-axis shows the number of markers used to calculate the genomic relationship matrix for G and G+P approaches. Six different set numbers (500, 1500, 5000, 10,000, 50,000, and 100,000) were randomly selected from an initial set of 200k to create the subsets of markers. During the calculation of the relationship matrix, the initial set was filtered down to 128,202, and the subsets used contained an average of 321, 957, 3,200, 6,382, 32,040, and 64,112 SNPs, respectively. For the PS model, 730 wavelengths were used from the data from average 2019 and 2020. The traits were evaluated in 2019, 2021, and 2022, including days to pollination (DTP), ear height (EH), ear length (EL), ear width (EW), germination (GER), plant height (PH), and tip fill (TPF).

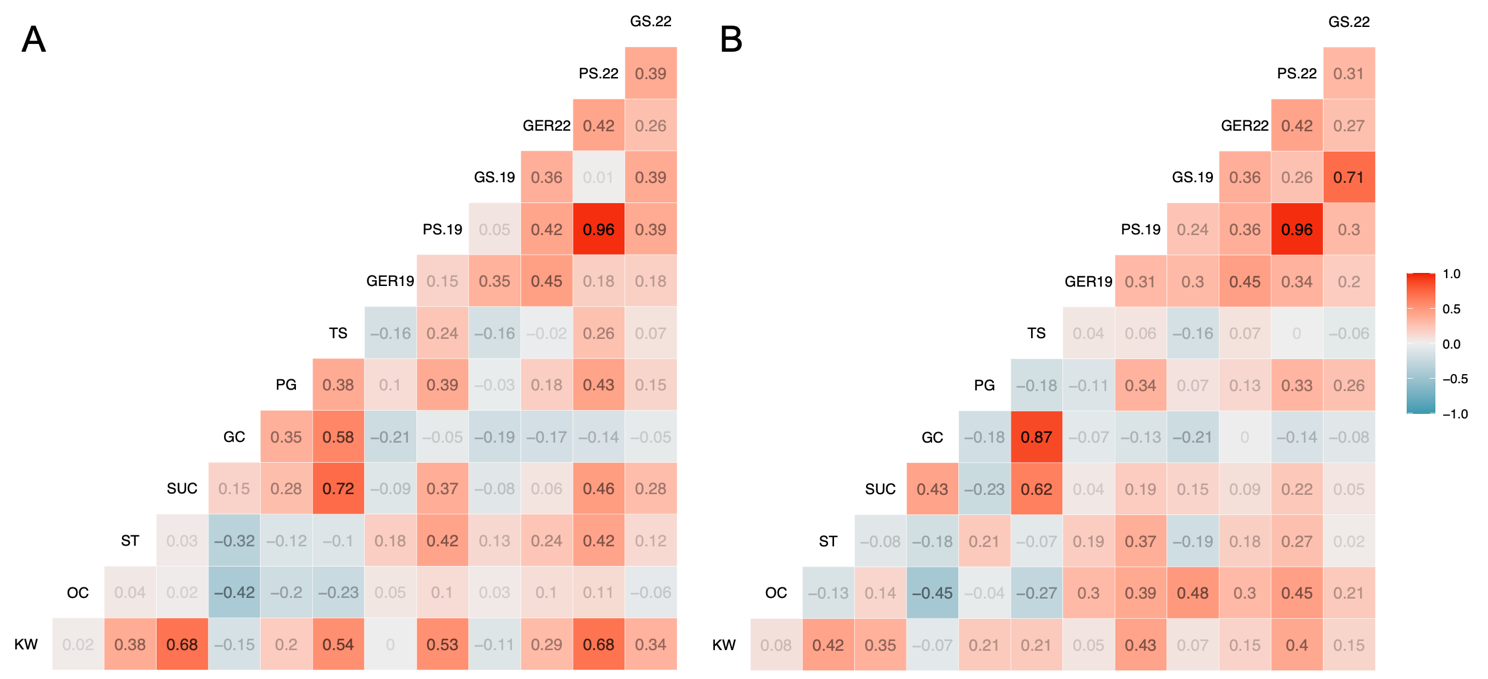

**Fig. S10** Correlation matrix displaying the relation between genomic and phenomic predictions with kernel composition traits. The kernel composition traits were measured using the NIRS with PLS method, which includes kernel weight (KW) (mg), oil content (OC,%), starch (ST,%), sucrose (SUC, mg), glucose (GC,%), phytoglycogen (PG, %), and total sugar (TS,mg). The GER represents the BLUEs of the germination rate observed in 2019 and 2022. The PS and GS represent the predicted values for the germination rate obtained from phenomic and genomic prediction (i.e. PBLUP and GBLUP), respectively. The predictions were made within groups in the diversity panel with mutations in the *sugary1* (A) and *shrunken2* (B) genes. Each cell in the matrix shows the Pearson correlation coefficient between pairs of values, with positive correlations displayed in red and negative correlations in blue. The PS model used the average NIRS data collected in 2019 and 2020 and the Standard Normal Variate first derivative preprocessing method.
